# Translation initiation by the Kozak mRNA sequence is based on a conformational readout on the ribosome

**DOI:** 10.1101/2025.07.11.664391

**Authors:** Ottilie von Loeffelholz, Charles Barchet, Samuel Holvec, Aida Abou Ramadan, Anne Maglott-Roth, S. Nimali T. de Silva, Isabelle Hazemann, Bruno P. Klaholz

## Abstract

The recognition mechanism of Kozak mRNA, typically comprising purines in the - 3 and +4 positions flanking the AUG start codon, has remained enigmatic for decades. To address this fundamental function in eukaryotes during translation initiation, we analysed the cryo-EM structures of human 48S preinitiation complexes with mRNA point mutations. They reveal a fan-like intercalation of the pre-codon triplet into the 18S ribosomal RNA (rRNA), while the -3 purine as opposed to the pyrimidine in the non-Kozak context favours stabilization of the ternary complex between initiation factor eIF2 and initiator tRNA. Specificity towards the +4 purine is achieved beyond a single residue recognition by mutual conformational adaptations of eIF1A, mRNA and rRNA and involves the insertion of a reading head in which decoding residue A1825 (rRNA) stacks with the A-site codon to stabilize the fully accommodated state. Hence, instead of relying on base pairing as in bacteria, the specific recognition of the Kozak sequence on eukaryotic ribosomes is based on an induced-fit mechanism that triggers a conformational readout of the mRNA.

## Introduction

Gene expression regulation at the translation level depends strongly on the messenger RNA (mRNA) sequence and secondary structure^1–4^. In bacteria, the correct positioning of the AUG start codon in the ribosome is favoured by the Shine-Dalgarno (SD) sequence that is found 8 nucleotides upstream of the AUG codon^5^; the SD sequence of the mRNA forms base-pairs with a complementary sequence at the 3’-end of the 16S ribosomal RNA (rRNA) located at the platform of the 30S body^6,7^, which allows to place the AUG start codon into the peptidyl (P) site. In contrast, in eukaryotes the AUG start codon is flanked by a different, eukaryote-specific nucleotide sequence called the Kozak sequence^4^, which was discovered by sequence comparisons of strongly translating genes and exists in many eukaryotic mRNAs^1,8,9^. The most conserved features of the Kozak sequence are a purine (A or G) at the -3 and +4 positions (preferably a G at the +4 position with the exception of fungi that have a bias for U^10^) relative to the AUG start codon (defined as +1, +2 and +3 nucleotide positions of the open reading frame, ORF). As a consequence, the most common start codon environment seen in eukaryotic mRNAs is (**A/G**)CC*AUG***G**, which is supposed to be recognized in some manner when the 48S pre-initiation complex (PIC) stops scanning the mRNA at the translation initiation site and forms the 48S PIC in which the initiator transfer RNA (tRNA_i_) sets the translation frame^11,12^.

Eukaryotic initiation factors (eIFs) play an important role during translation initiation and their general function has been elucidated in a number of structures of eukaryotic 48S PICs giving interesting insights into scanning, start codon recognition and subsequent steps towards subunit joining^13–19^. Specifically, initiation factors eIF2α and eIF1A are located next to mRNA nucleotides before and after the start codon, respectively^13,14,20–23;^ however, to understand how specific recognition of strong versus weak Kozak context is achieved and to address the role of individual nucleotides in a Kozak mRNA sequence would require comparisons with different nucleotides in the -3 and +4 positions, respectively. Only this would allow addressing the nucleotide selectivity of mRNA interactions involving eIF2α and eIF1A^13,14,20–23^, or components of the 40S small ribosomal subunit itself^24^, which are crucial to be understood at the single mRNA nucleotide level. In particular, the importance of Arg55 of eIF2α and Trp70 of eIF1A were speculated on previously^21,14,25^ but this region could neither be resolved well enough nor the experimental strategy enabled to clearly explain the meaning of the interactions regarding the strong preference for purines over pyrimidines in the Kozak mRNA.

We therefore set out to analyse the functional and structural determinants of mRNA point mutations guided by the efficiency differences that we measured using *in vitro* and *in vivo* translation assays. For this we structurally analysed and compared 48S PICs with Kozak and non-Kozak mRNA sequences, *i.e.* by switching purines into pyrimidines in the -3 and +4 positions, respectively. This analysis comprises also a 48S PIC in which eIF1A is mutated (W70A), which gives unique new insights into the selectivity role for efficient translation initiation of the mRNA versus this factor.

## Results

### Strong or weak Kozak sequences strongly differ in mRNA translation efficiency

We analysed human 48S PICs with Kozak and non-Kozak mRNAs functionally and structurally (**Fig. 1**). First, we reproduced Kozak’s previous observations^2,4^ in our human ribosome system using HeLa cell extracts with a model mRNA that contains the 5’ UTR of the human β-globin mRNA and coding for the firefly reporter (see methods; **Fig. 1A**). Our analysis reveals that (i) the best-preforming Kozak sequence tested contains an A-3 and a G+4 flanking the AUG start codon, (ii) substitution of the G in the +4 position by an A accounts for 50% translation efficiency loss, and (iii) additional replacement of the -3 purine (A) by a pyrimidine (C) or of the -2 and -1 positions from pyrimidine (C) to purine (A) further decreases the efficiency, with a pyrimidine stretch being the less efficient. Replacement of either sides of the AUG codon by uridines reduces the efficiency to below 20% (**Fig. 1A**). Based on these results, which are consistent with Kozak’s early work^4,26^, we designed a perfect Kozak and a non-Kozak sequence for further analysis of human 48S PICs, with variable nucleotides in the -3 or +4 nucleotide positions (ACC<u>AUG</u>GAA versus UUU<u>AUG</u>UUA). We then extended this analysis to the context of human cells to test the expression efficiency of vectors harbouring two reporter mRNAs comprising these two Kozak variants with the perfect Kozak sequence as internal control (see methods). The corresponding translation activity of these mRNAs differs significantly according to our fluorescence readout as observed for 2 different cell lines (HEK293T and hTERT, see methods and **Fig. 1B**) hence corroborating our mRNA sequence choice for the subsequent structural analysis.

**Fig. 1.**
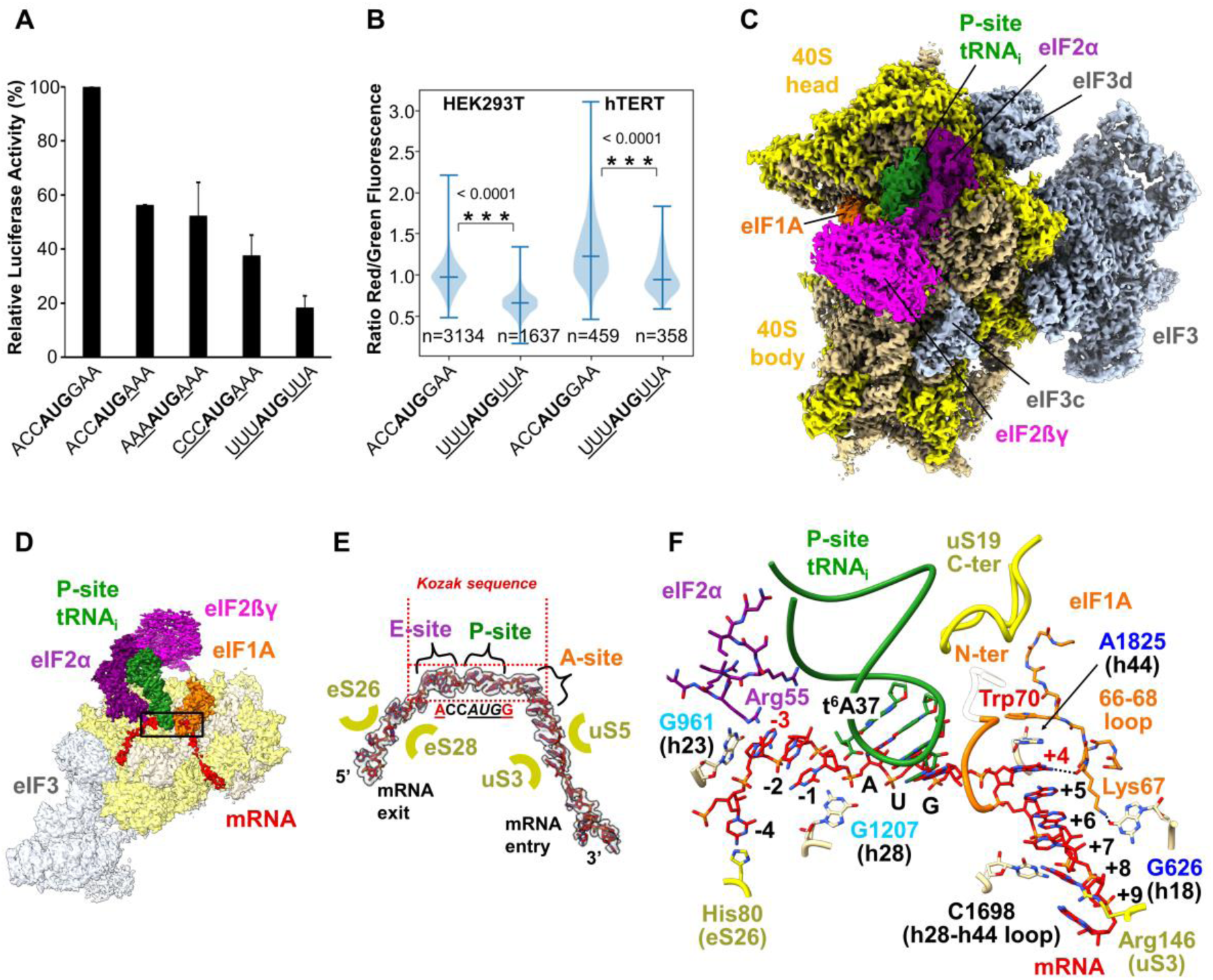
Functional and structural analysis of Kozak and non-Kozak 48S PICs. (**A**) Luciferase assay showing *in vitro* translation efficiency on mRNA harboring different Kozak mRNA sequences in HeLa cell extract. Shown are averages from 6 independent experiments together with their standard error of the mean. (**B**) *In vivo* translation assay using either HEK293T or hTERT cells 48h after infection. Shown is the violin plot with median and standard deviation of the relative fluorescence readout from translated Kozak (ACCAUGGAA) and non-Kozak (UUUAUGUUA) mRNAs *vs* Kozak mRNA (ACCAUGGAA, internal control, green). An unpaired t-test was applied revealing very significant translation differences of the start codon context in both cell lines (n: number of single cells imaged). (**C**) Cryo-EM structure of the human 48S PIC harboring the Kozak mRNA at 2.8 Å resolution; ribosomal proteins are colored yellow, rRNA ochre, mRNA red, tRNA_i_ green, eIF1A orange, eIF2α purple, eIF2βγ magenta and eIF3 light blue. (**D**) Top view of the structure shown in (C) using the same color code. (**E**) Depiction of the cryo-EM map density and atomic model of the Kozak mRNA (ACCAUGGAA) inside the human 48S PIC in the same orientation as (D). (**F**) Overview of the interactions formed by the mRNA Kozak sequence with the 40S small ribosomal subunit, tRNA_i_ and initiation factors eIF1A and eIF2α.

### Overall structure of the Kozak and non-Kozak mRNA 48S PICs and the environment of the Kozak sequence

For the comparative structural analysis of 48S PICs with Kozak and non-Kozak mRNA sequences we used the same *in vitro* translation system in HeLa cell extracts as for our functional analysis and purified the two complexes at the same time and under the same conditions (using GMPPNP to stabilize the eIF2-tRNA_i_ ternary complex, but no cross-linking; see methods). We implemented a mild complex isolation procedure from our *in vitro* translation assay under near-native conditions using cell extracts (see methods) thus bypassing the need for complete reconstitution. These complexes were analysed by high-resolution cryo-EM and advanced structure sorting techniques to address inherent sample heterogeneity^27–35^ (**Extended Data Fig. 1**). Using the perfect Kozak consensus sequence we obtained the structure of the corresponding human 48S PIC at an overall resolution of 2.8 Å (**Fig. 1C & D** and **Extended Data Figs. 1-3** as also illustrated from the local resolution side-chain features, see **Extended Data Figs. 4 & 5E**). This complex contains the ternary complex with tRNA_i_ and eIF2, eIF1A and eIF3 (**Fig. 1C & D**; factor ABCE1 was also seen as reported by others^21,36,37^ but not analysed further here, see **Extended Data Fig. 1**). The P-site tRNA is flanked by eIF2α in the E-site and by eIF1A in the A-site, respectively, similarly to previous structures^13,14,21,25,38,39^. Importantly, the side-chains of all key residues of the two eIFs are well-ordered (**Extended Data Figs. 4 & 6**). The mRNA locks the 40S ribosomal subunit in a “closed” conformation with tRNA_i_ bound and fully engaged in start codon recognition, which resembles the previously observed (P_IN_) state^16,40^ and exhibits interactions involving chemical modifications of tRNA (t_6_A37) and rRNA (m^1^acp^3^Ψ1248) around the codon-anticodon mini-helix (**Fig. 1D**; see methods). The mRNA path can be seen from its entry between helix h18 of the 18S rRNA and the beak of the 40S head until the exit (**Figs. 1D & E, Extended Data Fig. 5A**) and the -4 to +9 nucleotide region can be precisely fit (**Fig. 1F**). The coding sequence downstream of the AUG start codon is located in the A site (bases +4 to +6) where eIF1A is bound (**Extended Data Fig. 5B-D**). A remarkable number of π-stacked bases including mRNA nucleotides +4 to +7 as well as C1698 (h28-h44 loop) and decoding residue A1825 of helix h44 (18S rRNA) can be observed (**Fig. 1F**) indicating a high structural order of the mRNA is this translational state (**Extended Data Fig. 5C**). Decoding residues A1824 and A1825, two nucleotides that are involved in tRNA recognition (1492/1493 in the *Thermus thermophilus* ribosome^41^), are found in an intermediate flipped-out conformation (**Fig. 1F**). The structure of the 48S PIC harbouring the non-Kozak mRNA was obtained at a similar resolution (2.9 Å; **Extended Data Figs. 1-4**) and shows the same head conformation, but major conformational differences of mRNA nucleotides in the -3 and +4 regions and of eIF2α and eIF1A amino acids are visible, providing unprecedented insights into the specificity of interactions in the -3 and +4 positions, as described in the following.

### Upstream base triplet intercalation into a G-clamp of the 18S rRNA and Kozak sequence recognition of the -3 purine (not pyrimidine) by eIF2α

A zoom onto the mRNA at the A-, P- and E-sites of the Kozak mRNA 48S PIC shows a segmentation of the mRNA bases into base-triplets (**Fig. 1F & Fig. 2 B, D & F**). The - 3 to -1 mRNA nucleotides of the Kozak sequence are located on the 5’ side of the AUG start codon, close to the E-site (**Fig. 1F**), and insert as a base triplet into a cavity between nucleotides G961 and G1207 of helices h23 and h28 of the 18S rRNA, respectively, forming base and ribose stacking on either side (**Fig. 1F & 2B**). These nucleotides are positioned in an orthogonal manner to form a clamp, thus requiring the triplet bases to progressively adapt their angular orientation to form a fan-like intercalation (**Fig. 2D**). This mRNA recognition mechanism involves no base pairing and the triplet insertion requires no conformational changes of nearby nucleotides as compared to an empty human ribosome^30,34,42^, indicating that the G-clamp is pre-configured.

**Fig. 2.**
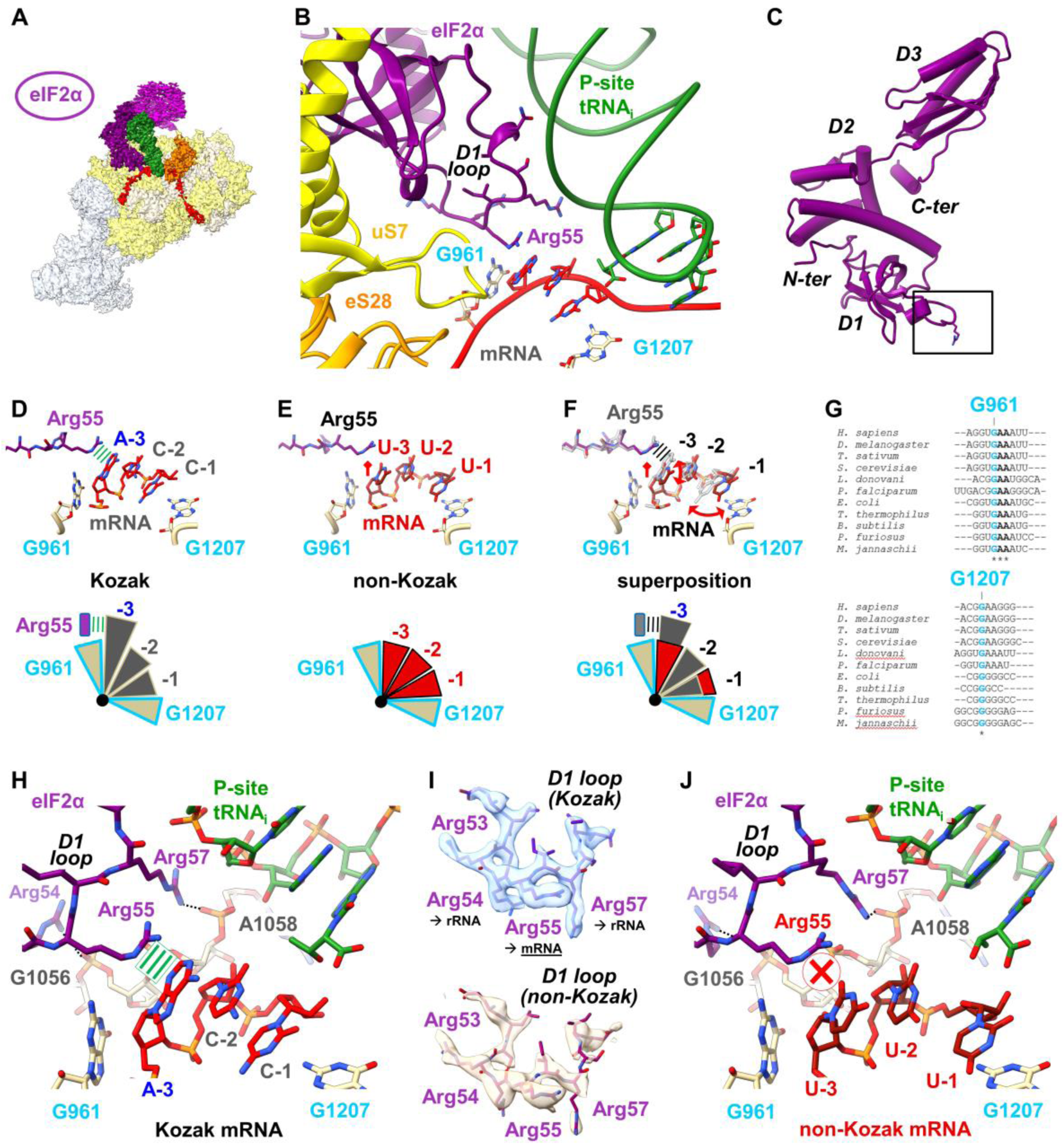
Interactions of the -3 to -1 nucleotide region of the Kozak and non-Kozak mRNAs with the preformed G-clamp pocket and eIF2α. **(A)** Overview of the cryo-EM structure of the human 48S PIC harboring the Kozak mRNA; eIF2α is highlighted in magenta. **(B)** Zoom on the upstream part of the Kozak sequence with respect to the AUG start codon showing the interaction environment of the -3 to -1 bases. **(C)** Overall structure of eIF2α. Domains 1, 2 and 3 (D1-3) as well as the N-terminus and the C-terminus are indicated. The location of the D1 loop that interacts with the -3 mRNA base is highlighted by a box. **(D)** Recognition of the upstream triplet of the Kozak mRNA by a preformed G-clamp (G961 and G1207, 18S rRNA) and eIF2α. Arg55 stabilizes the TC by interacting with A-3 of the Kozak mRNA via cation π-stacking. **(E)** Recognition of the upstream triplet of the non-Kozak mRNA by a preformed G-clamp but not by eIF2α that shows no interactions with the mRNA. **(F)** Superposition of (E) and (F). In comparison to the Kozak mRNA triplet, where the - 3 purine interacts with Arg55, the non-Kozak mRNA triplet appears retracted inside the G-clamp and shows no interactions with eIF2α at the level of the -3 pyrimidine, which is shorter. **(G)** Structure guided sequence alignment showing that the G-clamp is highly conserved. PDB codes used are: 6OKK, 6AZ1, 4V7E, 4V6W, 4V88 (18S rRNA) and 5L3P, 5IMQ, 5NJT, 4V4N, 4V6U (16S rRNA). **(H)** Overview of the D1 loop of eIF2α in the 48S PIC harbouring a perfect Kozak mRNA. Arg54 and Arg57 within the D1 loop undergo interactions with the phosphate backbone of G1056 and A1058 of the 18*S* rRNA, respectively. Arg55 forms cation π-stacking with A-3 of the Kozak mRNA. **(I)** Close-up view on the cryo-EM map and atomic model of the eIF2α D1 loop in the human 48S PIC harbouring a Kozak (top, blue) and non-Kozak (bottom, orange) mRNA. **(J)** Overview of the D1 loop of eIF2α in the 48S PIC harbouring a non-Kozak mRNA. The interactions of Arg54 and Arg57 within the D1 loop with the phosphate backbone of G1056 and A1058 of the ribosomal rRNA, respectively, are slightly altered in comparison to the 48S PIC on the Kozak mRNA. Arg55 forms no interactions with the mRNA here.

The -3 adenosine is found in close proximity to eIF2α at the level of the domain 1 (D1) loop, which reaches far into the E-site on the 40S ribosomal subunit to contact the mRNA (**Fig. 2A-C**), while the D1 & D2 domains of eIF2α interact with the P-site tRNA and ribosomal protein uS7 (**Fig. 2B & C**), respectively. The entire D1 loop has well-defined density in the cryo-EM map (**Fig. 2C, H & I** and **Extended Data Fig. 4 & 6**; better defined than in any previous structure^13,14,21,25,43^) and comprises a series of arginine residues (Arg53, Arg54, Arg55 and Arg57) among which only Arg55 contacts the mRNA (**Fig. 2D & H**). By contrast, the neighbouring Arg53 interacts with ribosomal protein uS11 (Lys63), while Arg54 and Arg57 interact with the phosphate backbone of the 1056 and 1057/1058 region of the 18S rRNA, respectively (**Fig. 2H & I**).

Arg55 forms a π-stacking interaction with the A-3 base of the Kozak sequence on the 5’ side of the G-clamp (**Fig. 2D & H** and **Extended Data Fig. 6E**). This cation-π interaction of the planar arginine guanidinium moiety with the aromatic -3 purine base is favoured by the delocalisation of the positive charge in the conjugated π-system and allows eIF2α to introduce a specific interaction with a purine in the -3 position. The observed stacking would also apply for a guanidine (G), because both purines (G or A) contain aromatic double-ring systems, but would not be favourable for pyrimidine nucleotides (C and U) as we show experimentally here next.

### A uridine in the -3 position leads to eIF2α specificity loss and triplet rearrangement in the G-clamp

The structure of the non-Kozak 48S PICs reveals that Arg55 of eIF2α does not interact with the -3 uridine base because it falls short to form a π-stacking interaction due to the smaller size of the single cyclic ring compared to the aromatic double-ring system of the purine in the Kozak complex (**Fig. 2E, F & J** and **Extended Data Fig. 6F**). Compared to purine, pyrimidine is only weakly aromatic due to the disruption of the electron delocalization by the keto substituent and the heterocyclic nature of the compound, which disfavours cation π-stacking. The interaction loss of Arg55 results in a partial destabilisation of eIF2α (D1 loop residues less resolved (**Fig. 2I**), thereby stabilizing the eIF2-tRNA_i_ ternary complex less strongly; **Extended Data Fig. 4AB & 6 E-H**; visible also in the globally refined Kozak versus non-Kozak maps, **Extended Data Fig. 1**) because the -3 uridine base can neither stack nor H-bond with the -3 base due to the unfavourable geometry. Due to these chemical properties, both U and C would not be recognized by Arg55, thus explaining the selectivity towards purines over pyrimidines in the -3 position the mRNA, clarifying its role as compared to previous structures where this side-chain was not clearly modelled^13,14,21,25,43^ (**Extended Data Fig. 7**).

The insertion of the UUU triplet into the G-clamp results in a more equally distributed stacking, which appears to be more extended to make the mRNA bridge the gap (∼15-20 Å) between the 40S head (G1207) and initiation factor eIF2α. An indirect effect of the Arg55 recognition loss for the -3 base is that the -1 base is also tilted towards the 3’ side of the G-clamp (**Fig. 2 D-G**), consistent with its broader EM density (**Extended Data Fig. 6E & F**) suggesting a more flexible positioning. Taken together, a pyrimidine in the -3 position cannot be recognized and therefore changes the interaction pattern at molecular level, consistent with the observed activity drop *in vitro* and *in vivo* compared to a -3 purine in the Kozak sequence (**Figs. 1A & B**).

### Kozak sequence recognition in the +4 position by rRNA nucleotide A1825 and eIF1A

Within the structure of the 48S PIC with the Kozak mRNA the +4 nucleotide is found to interact with decoding residue A1825 of the 18S rRNA (**Fig. 3A**). This residue forms two π-stacking interactions, on one side with the +4 purine base and on the other side with Trp70 of eIF1A, resulting in a characteristic triple-stacking involving 3 parts: mRNA, 18S rRNA and eIF1A protein (**Fig. 3A**). The direct stacking between A1825 and the +4 position is consistent with previous crosslinking experiments^20^, while in other cryo-EM studies of 48S PICs A1825 and Trp70 are swapped and oriented differently^14,21^ (as discussed below; see **Extended Data Fig. 8**). The A1825/+4 stacking extends further and also comprises nucleotide bases +5, +6 and +7, resulting in an extensive, long-range stacking (∼25 Å) in which the +4 nucleotide is embedded, hence not only recognizing the +4 purine but also forming a well-defined mRNA structure of the A-site codon and beyond (with the C1698 base intercalating between +7 and +8; **Fig. 3Ai**). In addition, eIF1A forms a hydrogen bond between the carbonyl peptide backbone of Lys67 and the 2-amino group of the G+4 base, which is the only specific interaction of eIF1A with G+4 (**Fig. 3Aii**).

**Fig. 3.**
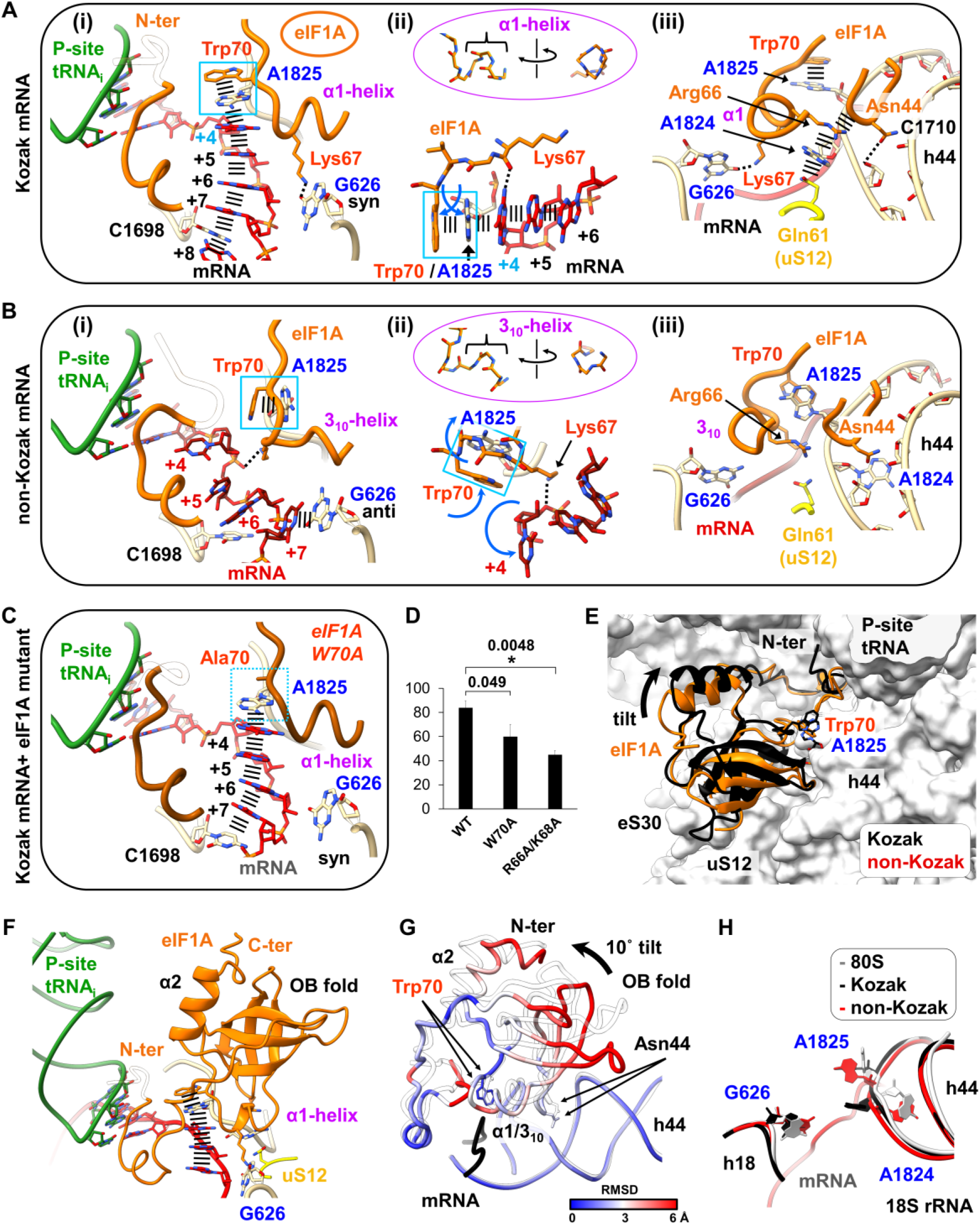
The mRNA sequence triggers an induced fit comprising mutual conformational adaptations of eIF1A, mRNA and rRNA in the Kozak 48S PIC compared to the non-Kozak 48S PIC. **(A) (i)** The Kozak mRNA 48S PIC shows extensive stacking of the A-site codon involving +4 purine recognition by decoding residue A1825, which in turn stacks with Trp70 of eIF1A. The α1 helix of eIF1A is indicated in magenta. **(ii)** Rotated view showing the specific interaction of G+4 through hydrogen bonding with the peptide backbone of Lys67 (eIF1A). **(iii)** View on h44 and loop L12 of eIF1A, showing eIF1A interactions with decoding residues G626, A1824 and A1825. **(B)** Structural rearrangements in the non-Kozak 48S PIC showing no +4 pyrimidine recognition by A1825. **(i-iii)** Close-up view as in (A). **(i)** The eIF1A residues 66-69 form a 3_10_ helix in this 48S PIC, Lys67 does not interact with decoding residue G636. **(ii)** The A1825/Trp70 duplex is retracted from the mRNA channel. **(C)** Structure of the of the Kozak mRNA 48S PIC with the eIF1A W70A mutant revealing the same conformation as seen in the 48S PIC with WT eIF1A (A). Decoding residue A1825 stacks with the +4 base even when Trp70 is mutated to Ala. Close-up view as in (A(i)) and (B(i)). **(D)** Luciferase assay showing *in vitro* translation efficiency on the Kozak mRNA in HeLa cell extracts which were supplemented with either wild type or mutant eIF1A. Shown are averages from 4 independent experiments together with their standard error of the mean as well as p-values calculated by an unpaired student’s t-test. **(E)** Interface view zoomed on the A-site showing conformational differences between the Kozak (black) and non-Kozak (orange) 48S PICs. **(F)** Overview of eIF1A and the mRNA in the A-site with the eIF1A N-terminus, OB fold, α1- and α2 helices and the C-terminus indicated. **(G)** Overlay of eIF1A of the Kozak 48S PIC (transparent white) and eIF1A of the non-Kozak 48S PIC (coloured by RMSD with respect to eIF1A in the Kozak 48S PIC). RMSD values are lower close to the mRNA and higher at distal parts of the factor. **(H)** comparison of the three decoding residues G262, A1824 and A1825 in the structures of empty 80S (light gray, PDB: 8QOI^34^), the Kozak 48S PIC (black) and the non-Kozak 48S PIC (red).

Compared to the Kozak complex, the structure of the non-Kozak mRNA 48S PIC reveals a major rearrangement of the mRNA (**Fig. 3B**). The triple-stacking is resolved, with A1825 (18S rRNA) and Trp70 (eIF1A) flipped aside, while the entire U+4 nucleoside (including the ribose moiety) appears rotated and shows no contacts (**Fig. 3Bii**). This is reminiscent of a lack of +4 pyrimidine recognition. In this conformation, A1825 and Trp70 also stack with each other as a duplex but are retracted from the mRNA channel (**Fig. 3Bii**). Hence, the non-Kozak mRNA is much less well-ordered and not stabilized, and decoding residue A1825 and Trp70 are found in an inactive conformation where the recognition of the +4 nucleotide cannot occur (**Fig. 3Bii**), consistent with the strongly reduced translational activity of a U+4 containing 48S PIC (**Fig. 1A & B**). The +4 interaction loss that originates from the strongly reduced aromaticity of pyrimidines compared to purines explains why both U and C are not recognized by A1825^44^. An additional factor contributing to fine-tuning the recognition mechanism is the H-bond interaction of the eIF1A peptide backbone with the 2-amino group of the G+4 base (**Fig. 3Aii**), which creates specificity towards G over A (where no such interaction is possible), consistent with the observed reduction of translational activity for an A (**Fig. 1A** and ref. 45).

The strongly differing conformations of the Kozak and non-Kozak complexes prompted us to analyse whether their conformations are unique. Further 3D classifications and refinement of the Kozak 48S PIC revealed that a second conformation exists (Kozak_2; see **Extended Data Fig. 8**) in which the extensive base stacking in the A-site is not formed, while Trp70 shows a peculiar stacking with the +4 base resembling that of previous structures of 43S and 48S PICs^14,21,36^. It is similar to the non-Kozak complex (which upon 3D classification shows only one dominant conformation, **Extended Data Fig. 1**) regarding the overall, non-stabilized mRNA conformation, but it differs in that the +4 nucleotide is reaching out of the mRNA channel to π-stack with Trp70 like in other 48S PICs^21^ (**Extended Data Fig. 8**). Structure comparisons show that the A1825/Trp70 duplex is retracted from the mRNA channel in both Kozak_2 and the non-Kozak complexes and in 3 previous studies^14,21,36^, while it is fully inserted only in the present Kozak_1 conformation where it is involved in an extensive, much more stabilized +4 base stacking in the A-site further consolidating the overall eIF1A and mRNA conformation (**Extended Data Fig. 8**). This stabilized, characteristic configuration is reminiscent of a specific +4 recognition. The occurrence of several conformational states highlights that an equilibrium exists between stabilized and non-stabilized complexes, which suggests that the mRNA undergoes a conformational sampling and sensing by the 48S PIC.

### A conserved sequence motif in eIF1A is responsible for the mRNA conformational readout

The 63-71 region of eIF1A contains a highly conserved GKLRKKVWI motif (comprising Lys67 and Trp70; **Extended Data Fig. 9**) of which the first part forms a 3_10_ helix (residues 63-66) and the second part forms a loop in the non-Kozak complex (**Fig. 3B**), whereas it adopts a classical α-helix (α1) in the Kozak mRNA complex (**Fig. 3A**). While Trp70 stacks with A1825, the neighbouring second decoding residue A1824 is stack-embedded in between Arg66 (eIF1A) and Gln61 of ribosomal protein uS12 (**Fig. 3Aiii**), resulting in π-stacking that stabilizes A1824 in a defined half flipped-out position in helix h44. This specific conformation is also stabilized by loop L12 of eIF1A, which reaches into the minor groove of helix h44 (18S rRNA) and interacts via Asn44 with the free 2’OH moiety of the A1823 ribose and with C1710 of helix h44 (**Fig. 3Aiii**). The specific A1824 conformation imposes a geometry on the phosphate backbone of the rRNA, which triggers decoding residue A1825 and Trp70 to move as a stacked duplex unit into the mRNA channel and interact with the G+4 Kozak base. Finally, the third decoding residue, G626 (18S rRNA) also changes conformation upon Kozak sequence recognition and retracts slightly (1 Å) from the mRNA channel, resulting in an H-bond between its 6-keto group and the side-chain of Lys67 (eIF1A; **Fig. 3Aiii**), a residue that is also specifically stabilizing the G+4 base through an H-bond with its carbonyl backbone (**Fig. 3Aii**).

In the non-Kozak complex the different tertiary structure arrangement of the conserved GKLRKKVWI motif towards a loop structure followed by a 3_10_ helix also repositions the Arg66 and Lys67 side-chains (**Figs. 3Bii & iii**), resulting in Lys67 interacting unspecifically with the mRNA phosphate backbone instead of with the +4 nucleotide, while Arg66 contacts the phosphate backbone of the 18S rRNA next to A1824 instead of stacking with its base (**Figs. 3Bii & iii**). Additionally, due to the overall OB domain positioning (**Fig. 3G**) Asn44 cannot reach as far into the minor groove of h44 as in the Kozak 48S PIC. As a result, decoding residue A1824 is found in an almost flipped-in conformation (**Fig. 3H**). Hence, the structural transition from a 3_10_ helix to a more stable α-helix conformation turns out to be a characteristic feature of the +4 Kozak recognition and contributes towards stabilization of the 48S PIC.

### G+4 recognition is based on an induced fit mechanism that triggers major conformational changes

The conformational transitions between Kozak and non-Kozak mRNA 48S PICs display a mutual adaptation of a series of residues around the +4 position, much beyond Trp70 of eIF1A alone, which comprise three components: the mRNA, key elements of the 18S rRNA and eIF1A. Major conformational changes are triggered by the G+4 readout, including the spring-like conformational transition of the 3_10_ helix to a classical α-helix and the formation of extensive stacking in response to recognition by A1825 (**Fig. 3A** versus **3B**). The positioning of the eIF1A N-terminal domain is similar in both complexes, while the globular part of the factor comprising the OB fold and the C-terminal helical domain move as an entity with the mRNA as pivot (**Fig. 3G**). As a result, the conformation and positioning of eIF1A differ significantly between the two complexes, with a tilt of the eIF1A OB fold by ∼10° towards the 40S head in the Kozak mRNA 48S PIC (**Figs. 3E & G**). Consequently, only in the Kozak 48S PIC the L12 loop of eIF1A (residue region 42-45, predicted to interact with RNA^46^) inserts far enough into the minor groove of helix h44 (18S rRNA) to induce an almost flipped-out conformation of A1824 and A1825 (**Figs. 3A, B & H**). This conformational adaptation complements the major rearrangement of the triple-stacking that involves A1825, the mRNA A-site codon and Trp70 of eIF1A and shows that the recognition of the +4 nucleotide of the Kozak mRNA relies on an induced fit mechanism.

Considering the structural involvement of eIF1A residues of the 66-68 loop (Arg66 stacking with A1824) and of Trp70 within the duplex stacking with A1825 we decided to functionally address their role for the recognition of the +4 mRNA position. Translational activity assays using the luciferase reporter system (see methods) show that cell extracts containing an excess of eIF1A W70A have a minor activity reduction (**Fig. 3D**). In contrast, altering the 66-68 loop (double mutant R66A/K68A) has a strong (statistically significant) effect (see methods), indicating that this eIF1A region is essential for its activity. As this is the central part around the region in which the largest conformational changes occur in eIF1A between our Kozak and non-Kozak 48S PICs, these residues appear to play a key role in triggering the induced fit mechanism.

### Structure of the 48S PIC complex with eIF1A W70A reveals that the mRNA sequence triggers +4 Kozak recognition

Based on the observations above and considering that Trp70 is the residue in direct stacking contact with decoding residue A1825 in the Kozak_1 complex or with the +4 purine in the Kozak_2 complex, we wondered which influence a mutation of this residue to alanine may have. We therefore set out to determine the cryo-EM structure of the 48S PIC Kozak mRNA complex in which eIF1A is replaced by eIF1A carrying the point mutation W70A (see methods). It reveals that the overall conformations of the Kozak mRNA and of eIF1A are the same (**Fig. 3C**). Importantly, A1825 maintains the direct stacking with the +4 purine also in absence of the Trp70 side-chain (**Fig. 3C**). The comparison with the WT eIF1A 48S PIC (**Fig. 3Ai**) shows that Trp70 certainly helps to stabilize the complex through stacking with A1825, but it does not *per se* induce the +4 purine-recognizing conformation of mRNA, rRNA and eIF1A. Instead, the mRNA sequence itself appears to be the main driver, because a single pyrimidine in the +4 position abrogates the formation of an mRNA structure that can be recognized (**Fig. 3B**). This also shows that among the two residues involved in the stacking duplex, it is the rRNA component (A1825) that triggers +4 recognition. In accordance with the structural data of the 48S PIC with eIF1A-W70A no significant defects in *in vitro* translation were observed when an excess of the mutant was added in comparison to WT eIF1A (**Fig. 3D**). Taken together, the induced fit mechanism seen for the Kozak mRNA 48S PIC with WT eIF1A occurs in the same manner in the W70A mutant complex.

## Discussion

In this study we assessed the translational activity of Kozak and non-Kozak mRNA sequences harbouring mutations in the -3 and +4 nucleotide positions and determined the structure of the corresponding 48S PICs, which allowed to directly compare the response of 48S PICs towards different start codon environments. While the role of eIF1A Trp70 was speculated on previously^14,21^, more residues (in particular Arg66) were proposed based on the NMR structure of the isolated factor^46^, but their role has remained unclear. The present mutational analysis in the context of Kozak and non-Kozak mRNA sequences and the structural analysis of the corresponding 48S PICs, including that of the W70A mutant, provide experimental evidence for the molecular basis underlying Kozak selectivity. They clarify that +4 base recognition is based on a series of residues, extending much beyond Trp70, comprising a whole region of eIF1A and involving a mutual conformational adaptation of eIF1A, mRNA and rRNA. Accordingly, the neighbouring loop residues (Arg66/Lys68) turn out to play a previously non-anticipated important functional role. Moreover, our results demonstrate how the sequence difference in the two mRNAs is directly linked to substantially different translation rates and structural readouts where the mRNA adopts different conformations inside the mRNA channel of 48S PICs. We show that the conformational selection of the mRNA triggers structural rearrangements in the 18S rRNA of the 40S ribosomal subunit as well as in initiation factors eIF2α and eIF1A, demonstrating that the recognition of the Kozak mRNA relies on an induced-fit mechanism^47^.

With regards to the role of the ribosome, the structures clarify that the Kozak sequence actually *does* interact with some key components of the rRNA, but instead of the complementary base *pairing* through the formation of an SD/anti-SD duplex seen in bacteria^5–7^, in eukaryotes this interaction relies on base *stacking* involving key elements of the 18S rRNA. Thus, the primary environment is rRNA-based, while eIF2α and eIF1A serve to introduce additional selectivity over the general accommodation of the mRNA. Interestingly, in both Kozak positions (-3 and +4), the nucleotides are stabilized as entire base triplets and it is the first nucleotide of each triplet that provides specificity. The base stacking turns out though to occur differently in the two positions (**Fig. 4**): in the -3 position, it comprises the insertion of the pre-codon triplet upstream of the AUG into a dedicated cavity in the 18S rRNA (the G-clamp) and the A-3 nucleotide base is directly recognized by Arg55 (eIF2α; **Fig. 2**) which in turn stabilizes the eIF2-tRNA_i_ ternary complex. The importance of the cation π-stacking now becomes clear from the systematic comparison of Kozak and non-Kozak complexes showing that Arg55 of eIF2α forms this interaction specifically only in the presence of a purine, while it does not in the presence of a pyrimidine. Regarding the +4 position, in the fully accommodated state (Kozak_1) the G+4 nucleotide is recognized by decoding residue A1825 of the 18S rRNA and indirectly by eIF1A (**Fig. 3**) through a series of interactions and conformational adaptations (discussed below). Hence, instead of a simplistic, direct readout of the -3 and +4 nucleotides, numerous components are involved leading to translation activation upon the induced-fit recognition (**Fig. 4**). Hence, an allosteric effect between the -3 and +4 regions could be plausible, which would create cross-talk and synergy between these regulatory sites to fine-tune expression levels^48^, consistent with the existence of a family of Kozak sequences rather than a “consensus” sequence.

**Fig. 4.**
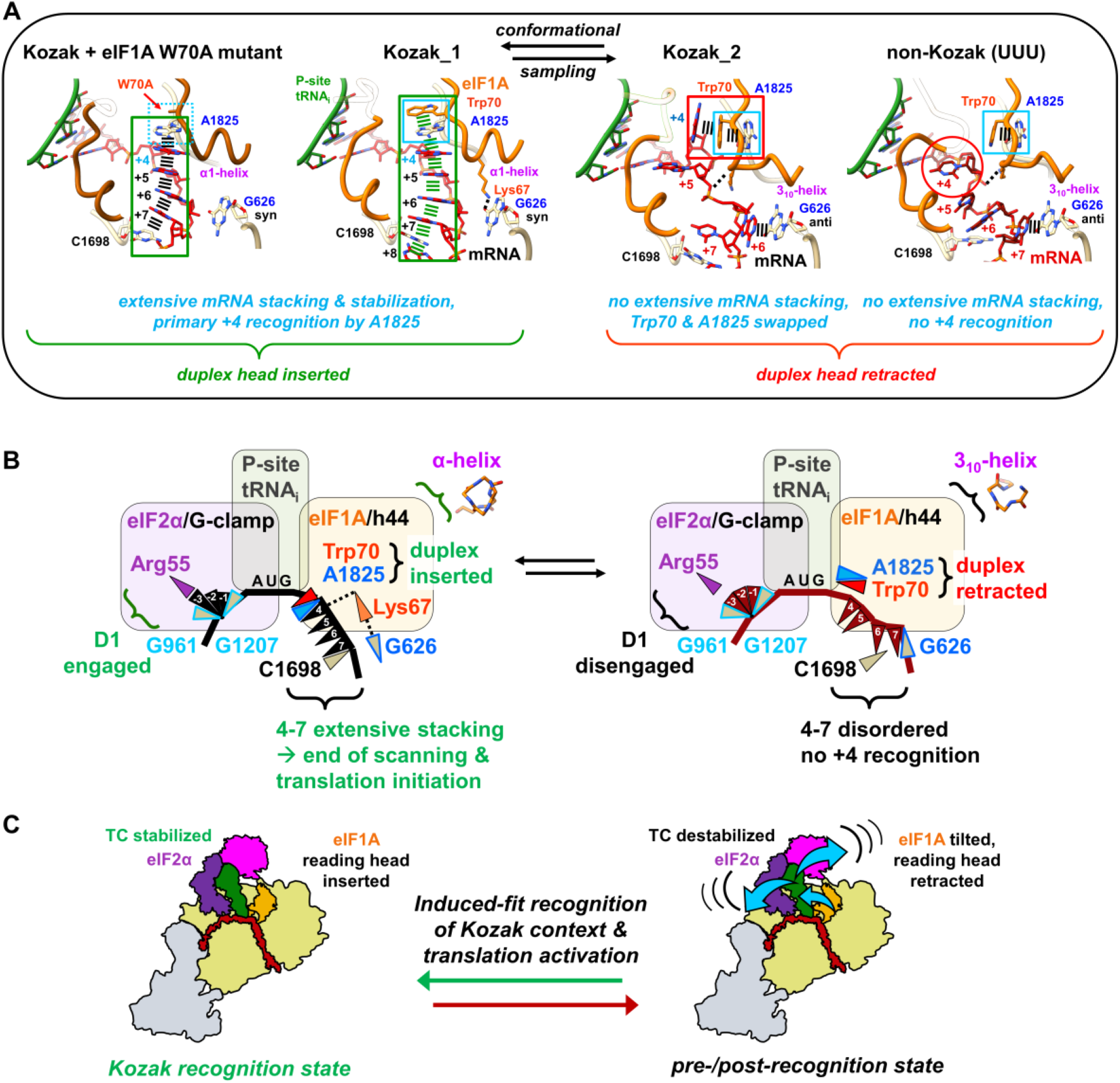
Schematic depiction of Kozak sequence selection by the 48S PIC during translation initiation. 48S PIC formation on the mRNA start codon after mRNA scanning can lead to two potential complexes. The presence of a Kozak context in the mRNA favors TC stabilization and translation activation through an induced fit mechanism that involves major conformational changes and a mutual adaptation of mRNA, rRNA and eIF2α and eIF1A, which triggers a conformational readout of the mRNA sequence on eukaryotic ribosomes. The activated state (left) exhibits an engaged eIF2α D1 loop interacting with the -3 purine, whereas eIF1A forms a more stable α-helix and the A1825/Trp70 duplex is inserted and recognizes the +4 purine leading to extensive stacking and stabilization of the A-site codon region, which also involves C1698 and decoding residue G626 of the rRNA. The resulting TC stabilization favors end of scanning and translation initiation. In contrast, the destabilized state (right) comprises a disengaged eIF2α D1 loop in the presence of a -3 pyrimidine and the mRNA inserts only into the G-clamp, while in the A-site the A1825/Trp70 duplex is retracted from the mRNA channel and does not recognize a +4 pyrimidine; eIF1A forms a short 3_10_ helix and the A-site codon is not well ordered.

The induced fit applies both for the -3 and the +4 purines, but with different configurations. In the -3 position, the triplet inserts into the pre-configured G-clamp that is specially shaped to fit a base triplet in the E-site, similar to a key-lock mechanism^49^, but with an additional induced-fit that accounts for the flexibility and adaptability of the eIF2α D1 loop and of the -3 purine versus pyrimidine in the mRNA, which the G-clamp forces into a segmented base triplet through a triple-base intercalation. The G-clamp pocket itself and Arg55 of eIF2α are providing a “lock-and-key” environment for the -3 nucleotide without significant conformational changes. The fan-like triplet stabilisation sets the frame with the ORF, thus positioning the AUG start codon into the P-site for the first decoding event to take place with the tRNA_i_, which in turn stabilises the head of the 40S ribosomal subunit in the P_IN_ state^16^. In eukaryotes, an interaction of the -3 base with G961 had been proposed based on a low-resolution yeast 80S ribosome structure^50^, which is now clearly resolved in the present study. Interestingly, the fan-like triplet insertion also exists in bacterial 70S ribosomes^51,52^, but due to the lack of eIF2 in eubacteria there is no specific recognition of the upstream region, explaining how the -3 Kozak context recognition became an evolutionary acquisition of eukaryotes.

The G-clamp triplet insertion *per se* is sequence-independent with regards to the mRNA. To introduce specificity towards the -3 purine hence requires Arg55 (eIF2α), where the cation-π stacking interaction allows recognition of both A and G. In contrast to previous assumptions, the neighbouring Arg53, Arg54 and Arg57 residues do not interact with the mRNA but instead with the 18S rRNA of the 40S ribosomal subunit and thus play an indirect role in maintaining the structure of the D1 loop and tRNA_i_ (**Figs. 2H & J**). While earlier observations suggested an involvement of both Arg55 and Arg57 based on proximity^25,38^ the present work clarifies that among the 4 adjacent arginine residues in the D1 loop only Arg55 is involved in Kozak base recognition. The overall positioning of eIF2α and Arg55 is similar to that seen in a recent study^14^ (see **Extended Data Fig. 7**) but the present data show a better accommodation of the -3 base, which is visible from the improved density (i.e. stabilization) of the D1 loop and better geometry of the π-stacking with the Arg55 side chain. Further, using a -3 mRNA mutant now clarifies the selectivity role of the -3 purine as opposed to a pyrimidine in the non-Kozak 48S PIC, the latter not being able to form cation π-stacking with Arg55 and resulting in reduced stabilization of the eIF2α D1-loop and as a consequence the ternary complex. As a result, the tRNA shoulder and acceptor arm are slightly tilted in the non-Kozak complex and may not be able to proceed efficiently towards ribosomal subunit joining and translation initiation (**Fig. 4** & **Extended Data Fig. 6G&H**).

For the +4 position, the induced-fit mechanism is more complex and involves mutual conformational adaptations of eIF1A, mRNA and rRNA, much beyond a simple recognition by a single residue (Trp70) assumed in previous studies, which also explains the mild effect of a single W70A mutant **(Fig. 3C&D)**. It comprises a conformational readout by the stacking duplex formed by decoding residue A1825 (18S rRNA) and Trp70 (eIF1A) accompanied by a secondary structure transition from a 3_10_- to an α-helix of eIF1A (**Fig. 3** and 4A). The key element turns out to be decoding residue A1825 of the 18S rRNA that primes +4 purine recognition by stacking with the G+4 base (equally possible with an A), thereby mediating stacking with Trp70 of eIF1A, while Lys67 provides additional fine-tuning specificity for G over A through H-bonding. Compared to previously described 48S PICs^14,21,43^, the position of Trp70 and A1825 are swapped, as discussed below. The structure of the 48S PIC with eIF1A-W70A underlines the central role of A1825, because in the absence of the Trp70 side-chain the same interactions and conformations take place between A1825 and the +4 purine. Yet, Trp70 accompanies the A1825 rotation and mRNA accommodation upon Kozak mRNA recognition as compared to the completely differently arranged non-Kozak mRNA (**Figs. 3** and **4A**), indicating only a complementary role of Trp70 in complex stabilizing. Rather than acting through a single residue-specific recognition of the +4 purine base, eIF1A triggers a conformational selection of the mRNA and of the A1825 conformation, *i.e.* the molecular recognition of the Kozak context in the +4 position is driven by the mRNA sequence. This concept is consistent with our observation that the reduction of the translational activity for 48S PICs containing eIF1A-W70A is not significant and that Trp70 mutation may reduce but not abolish 48S complex formation^46^. In contrast, mutating the region next to the 3_10_ helix (R66A/K68A double mutant) results in a significant activity loss. The presence of a 3_10_ helix had been actually noticed in the NMR structure of eIF1A^46^ (similar to that in the non-Kozak complex), the role of which now becomes clear in the Kozak context where it acts as a sensing “spring” that snaps in upon Kozak mRNA recognition and activation as part of the induced fit mechanism (**Fig. 4B&C**, left). By contrast, a +4 pyrimidine does not interact with the A1825 decoding residue and hence triggers no folding of the α1 helix, which results in an unlocked eIF1A conformation and an unproductive 48S PIC (**Fig. 4B&C**, right).

This study provides explicit evidence that 2 conformations of the Kozak complex exist comprising completely different interaction patterns at the +4 position. Kozak_2 is similar to previous structures^14,21,43^ (albeit better resolved now), while Kozak_1 shows a swapping of A1825 and Trp70, resulting in direct stacking of A1825 with the +4 purine and very extensive stacking of the A-site codon, i.e. forming a particularly stable complex. This implies that Kozak_1 is actually the activated state in which Kozak mRNA sequence recognition happens. Our data allow to integrate and rationalize previous data into a comprehensive mechanism, which is based on an mRNA-driven conformational selection (**Fig. 4**). Kozak_1 adopts a characteristic configuration in which A1825 recognizes the +4 purine, while the Kozak_2 conformation represents a non-recognizing, probably mRNA sensing complex, in which the mRNA is not well ordered & less stabilized and the A1825/Trp70 reading head retracted (**Fig. 4A**; like in previously described 48S PICs, which may represent other functional pre-initiation intermediates in equilibrium with each other^21,43,14^; **Extended Data Fig. 8**). In the non-Kozak complex, the mRNA is also less ordered, but due to absence of +4 recognition the pyrimidine is flipped away (**Fig. 4A**). Interestingly, before mRNA binding in the 43S PIC^36^ the A1825/Trp70 duplex is preconfigured to act as a reading head for a +4 purine, very much in the same position as in the Kozak_2 and non-Kozak complexes **(Extended Data Fig 7**), apparently waiting for the recognition event to happen and induce a conformational accommodation of the mRNA, as observed in Kozak_1 (**Fig. 3A**). In the non-Kozak 48S PIC the duplex does not interact with the +4 nucleotide (it is flipped away; **Fig. 3B**); the complex instead assumes a non-productive conformation in which the mRNA adopts an intermediate, less-ordered and non-stacked configuration, with the reading head in a retracted position (**Fig. 4A**), resembling that of recent studies with and without mRNA^36,21,14^. Taken together, the A1825/Trp70 reading head adopts either *inserted* or *retracted* conformations.

The activating and non-activating conformational states (**Fig. 4A**) are probably in equilibrium with each other and not incompatible *per se*, consistent with the existence of two conformations for the Kozak complex (Kozak_1/2), suggesting that a conformal sampling of the mRNA is occurring; in contrast, only one conformation is found for the non-Kozak complex. The present Kozak 48S PIC shows many novel features explaining specific translation activation, in particular a strong stabilization of the complex including extensive stacking of the A-site codon, pre-configured for subsequent tRNA binding during the elongation phase. The Kozak_1 conformation is actually consistent with and explains the crosslinking activity of the +4 position with A1824/A1825 found many years ago^20^, hence corroborating that the interaction of the +4 purine with these decoding residues is responsible for recognizing the +4 position. Finally, the structure of the eIF1A-W70A mutant complex shows that Trp70 does not *per se* induce the A1825 configuration, underlining that larger parts of eIF1A are involved in the recognition process, as illustrated by the fact that eIF1A is involved in the stabilization of all 3 decoding residues of the 18S rRNA (G626, A1824, A1825; **Fig. 3Aiii**). Taken together, this comprehensive structure-function study elucidates the molecular basis of mRNA Kozak sequence recognition on the ribosome and rationalizes the specific function of -3 and +4 nucleotides during translation initiation (**Fig. 4**). The base pairing of the SD sequence observed in bacteria is compensated by the specific recognition of the Kozak sequence in eukaryotes through an induced fit mechanism, resulting in a conformational selection of the mRNA that leads to stop of scanning, preparation of eIF5B-promoted subunit joining^14,53^ and transition to the elongation phase. In conclusion, this study addresses a fundamental mechanism in eukaryotes.

## Acknowledgments

We thank Jonathan Michalon, Mathieu Schaeffer and Nicolas Ballet for IT support, Léo Fréchin, Brice Beinsteiner & Charles Barchet for maintaining computing clusters, and Paola Rosolino and Nicole Jung from the CBI cloning platform for their support, the IGBMC cell culture facilities for HeLa cell production, members of the integrated structural biology platform at CBI for support, Rafael Schoch for help with Python scripting, Eric Huntzinger and Bertrand Séraphin for help with a luminometer, Albert Weixlbaumer for sharing his plasmid encoding the T7 RNA polymerase and for the purification protocol, and Adam Ben-Shem for fruitful discussions. This work was supported by CNRS, Association pour la Recherche sur le Cancer (ARC), Institut National du Cancer (INCa), the Fondation pour la Recherche Médicale (FRM), Ligue nationale contre le cancer (Ligue) and Agence National pour la Recherche (ANR), USIAS of the University of Strasbourg (USIAS-2018-012) and ANR JCJC (ANR-23-CE11-0003-01 Kozak to OvL). This work of the Interdisciplinary Thematic Institute IMCBio, as part of the ITI 2021-2028 program of the University of Strasbourg, CNRS and Inserm, was supported by IdEx Unistra (ANR-10-IDEX-0002) and by SFRI-STRAT’US project (ANR 20-SFRI-0012), EUR IMCBio (ANR-17-EURE-0023) under the framework of the France 2030 program and LabexNetRNA (ANR-10-LABX-0036_NETRNA) administered by ANR. The electron microscope facility was supported by the Region Grand Est, FEDER, the French Infrastructure for Integrated Structural Biology (FRISBI) ANR-10-INBS-0005 / France 2030 program, EquipEx^+^ France-Cryo-EM (ANR-21-ESRE-0046) and Instruct-ERIC.

## Author Contributions

O.vL., I.H. & S.N.T.D. and S.H. performed sample preparation, O.vL., S.N.T.D., I.H., A.A.R. & A.M-R. mutational analysis, O.vL. cryo-EM data collection & processing, O.vL & C. B. model building & refinement, O.vL. & B.P.K. structural analysis. All authors analysed the data. O.vL. and B.P.K. conceived the project and wrote the manuscript with input from all authors.

## Materials and Methods

### Vector plasmids

The β-globin 5’-UTR was cloned in front of the firefly luciferase gene in a pUC16 vector and variants were obtained by modifying the sequence next to the AUG start codon to ACC*aug*GAA, ACC*aug*AAA, AAA*aug*AAA, CCC*aug*AAA and UUU*aug*UUA (not UUU*aug*UAA to avoid creating a stop codon) using site-directed mutagenesis. The genes including their 5’-UTRs were amplified by PCR for further use. For *in vivo* studies the ACC*aug*GAA Kozak sequences was designed with a minimal eEF1A promoter followed by the β-globin 5’-UTR cloned in front of the TdTomato and Clover3 ORFs, which were set in tandem in the same plasmid separated by a polyA signal, so that both mRNA constructs ensuring transfection of all cells with red and green reporter mRNAs are independently transcribed from the same plasmid. A second plasmid was generated by mutating the ACC*aug*GAA Kozak sequence in front of TdTomato to UUU*aug*UUA. The clone of human eIF1A in the pET28a(+) vector was purchased from ProteoGenix. The eIF1A R66A/K68A and W70A mutants were generated by site-directed mutagenesis of this plasmid.

### *In vitro* transcription and mRNA constructs

We designed two fifty-nucleotides long mRNA sequences derived from the human β-globin gene to avoid using short synthetic mRNAs. The mRNA constructs comprise the β-globin gene with 20 nucleotides upstream including its natural Kozak sequence, the AUG start codon followed by a lysine codon (AAA) and a stop codon (UAA) and 21 nucleotides that occur in the natural β-globin mRNA ORF. This housekeeping gene is often used as a reference in PCR experiments, has no secondary structure and generates high protein expression levels. The mRNA constructs containing the gene of firefly luciferase with varying context around the AUG start codon were transcribed *in vitro* using 10 μg of PCR product, 7 mM NTPs, 0.8 U/μl RNAsine, 200 mM HEPES-KOH (pH 7.5), 30 mM Mg(OAc)_2_, 2 mM spermidine, 40 μM DTT and 2 U/μl of purified T7 RNA polymerase^54^. The *in vitro* transcribed mRNA was purified in two steps by LiCl precipitation and ethanol precipitation, respectively, and then capped and Poly-A tailed (enzymes from NEB).

Additionally, synthetic mRNA constructs describing short versions of the β-globin mRNA carrying the sequence 5’agcaaccucaaacagacNNN*aug*NNauaacugacuccugaggagaagucu3’ were ordered from Dharmacon (this construct has no secondary structure according to RNA folding predictions software^55^). Here the sequence NNNaugNNN was varied to ACC*aug*GAA, ACC*aug*AAA, CCC*aug*AAA and UUU*aug*UUA.

### *In vitro* translation and luciferase activity measurements

*In vitro* translation was done with 40% HeLa cell extract (Ipracell) as described in^56^ using optimized potassium acetate and magnesium acetate concentrations (**Extended Data Fig. 10**). Reactions were carried out with 100 ng mRNA in 20 μl sample volume at 37° C for 30 min. The *in vitro* translation was stopped by addition of 2x reaction lysis buffer (Promega) and luciferase activity was tested with a Luminometer (Berthold Technologies) with the firefly luciferase assay kit from Promega according to the manufacturer’s instructions. Luminescence for each *in vitro* translation was measured in triplicates with 50 μl of substrate solution with a read-out time of 10 s, and the experiment was repeated six times. The eIF1A mutant activity measurements were done similarly, only that eIF1A wild type or mutants were added in excess (0.08 mg/ml final concentration) to the translation mix. The experiment with excess eIF1A was repeated four times. Students t-test analysis was applied to test for readout significance.

### *In vivo* translation activity measurements

The reporter plasmids were transfected into HEK293T and hTERT cells. The cells were fixed by formaldehyde 48 hours after transfection and stained with DAPI. Images were acquired on a Cellomics Cx7 automated microscope at 10x magnification. Using HCSStudio software and TargetActivation Bioapplication, cells were identified using DAPI staining, and the nuclei mask was dilated to measure green and red intensity. Data were analysed using Python. For this, ratios of red (AUG start codon in a Kozak or non-Kozak context) vs green (AUG start codon in a Kozak as an internal reference) readout of each cell was plotted. Significance of the difference in mean ratios was computed using unpaired student’s t-test.

### Sample preparation

Eukaryotic 48S PICs of the human ribosome were isolated under near-native conditions using HeLa cell extracts via size fractionation of the translation mixture followed by cryo-EM analysis of the 40S fractions in which different conformations and compositions were sorted out in 3D by extensive focused classifications, signal subtractions and focused refinements^27–29,31^ to address inherent sample heterogeneity. This powerful combination provides a significant advancement in our ability to study these complexes in their native context. An excess of 50-nucleotide mRNA was used (5 µM) to compete out endogenous mRNAs. To inhibit factor dissociation and stabilize the complexes we added 1 mM non-hydrolysable nucleotide analogue GMPPNP (guanosine 5′-[β,γ-imido]triphosphate, Sigma) and performed size fractionation of the translation mixture using 15-30% sucrose gradient centrifugation from which the 40S fraction was subjected to negative-staining EM and subsequent cryo-EM. The 48S PICs with initiation factors and ACCAUGG mRNA or UUUAUGU mRNA were assembled and cryo-EM grids prepared at the same time. The fractions containing 48S PICs were pooled, concentrated by ultracentrifugation and resuspended in gradient buffer without sucrose containing 1 mM GMPPNP. Recombinant eIF1A WT and mutant were purified as previously described^57^. The endogenously isolated complexes were supplemented with additional eIF1A (∼100-fold excess) to achieve better stoichiometric binding as it tends to dissociate (this is a standard procedure to favour stoichiometric binding; shifting the equilibrium does not influence factor conformations *per se*, only their populations). No cross-linking was done. The sample was applied to plasma-cleaned (90% Argon, 10% oxygen; Fischione) holy R 2/2 carbon 300 mesh gold grids (Quantifoil) coated with a thin carbon film, blotted and plunge-frozen at 10°C and 95% humidity in liquid ethane using a Vitrobot IV (Thermo Fisher Scientific, The Netherlands).

### Cryo-EM data collection and image processing

42426 movies and 24852 movies for the 48S PICs with the ACCAUGG and UUUAUGU mRNAs were respectively collected on the in-house G4-Titan Krios equipped with cold-FEG, Selectris X energy filter and Falcon4i direct electron camera in counting mode using SerialEM^58^ in fringe-free alignment settings. The data collection was set to nine exposures per hole and set to image 9 holes per stage position, a pixel size of 0.72 Å at a dose-rate of ∼10 e^-^/pixel per second and a total dose of 40 e^-^/Å^2^ using the SerialEM software^58^ with a target defocus range of -0.5 μm to -2.5 μm. For the 48S PICs containing the ACCAUGG mRNA and the eIF1A W70A mutant 14280 movies were collected in the same magnification and dose settings on the same microscope using AFIS in the EPU software. Optics groups were assigned to the movies according to the beam tilt applied during microscope exposure to correct for off-axis coma^59^.

Data pre-processing of the EER movies was done using the software MotionCor2 v1.6.3^60^ for movie alignment and CtfFIND^61^ for correction of the contrast transfer function (CTF). Particles were selected with the software Gautomatch^62^ or RELION v3.0^63^ and the picked particles were cleaned by 2D classification in RELION v3.0^63^ or cryoSPARC^64^. Particle sorting by 3D classification as well as further 3D classifications, 3D refinements and map sharpening were performed in RELION v3.0^63^ including focused classifications and refinements of the 40S body and head parts and the tRNA, and CTF refinement^29–31,33^.

After 2D classification 287121 good particles of the Kozak 48S PICs harboring the ACCAUGAA mRNA were extracted with a down sampling to 4.32 Å/pixels. Due to the sensitivity of the 40S head conformation with respect to the presence of mRNA, tRNA and factors we could use this feature for 3D structure sorting (**Extended Data Fig. 1**). Focused classification on the head of the 40S small ribosomal subunit resulted in 121386 particles containing the 40S small ribosomal subunits in an open conformation, 61554 particles with a 40S small ribosomal subunit in a half-closed conformation and 104181 particles with 48S PICs in a closed head conformation. The particles with PICs were further classified focused on the ternary complex resulting in 19645 particles without tRNA_i_ or eIF2 and 84540 particles with tRNA_i_ and eIF2. The particles with tRNA_i_ and eIF2 were re-extracted and coarsened to 1.44 Å/pixels and the structure refined to 2.9 Å average resolution (**Extended Data Fig. 1-3**). Because we detected further heterogeneity in this map we decided to include a further 3D classification step for the 40S head that helped to differentiate three main states of 48S complexes containing 39433, 36575 and 8532 particles, respectively, which differed slightly in their head swivel or rotation (**Extended Data Fig. 1**). The largest population with 39433 particles was refined to a map at 3.0 Å resolution, which showed a well-defined mRNA conformation that was that was refined further by multibody refinement^33,65^ resulting in three maps with an overall resolution of 2.8 Å for the 40S body including mRNA and eIF1A, 2.8 Å for the 40S head together with the ternary complex and 4.2 Å for eIF3 and ABCE1 (**Extended Data Fig. 1**). The 40S body, 40S head & TC and eIF3 maps after partial signal subtraction with centering and focused refinement reached a resolution of 2.8 Å, 2.8 Å and 3.8 Å (**Extended Data Figs. 1-3**), respectively, and were amplitude-corrected using the auto_sharpen module of the Phenix software^66^, combined to a composite map using the software CHIMERA, and used for atomic model building and map interpretations.

After 2D classification 320378 good particles of the non-Kozak 48S PICs harbouring the UUUAUGUUA mRNA were extracted and coarsened to 4.32 Å/pixels. Focused classification on the head of the 40S small ribosomal subunit resulted in 176161 particles containing the 40S small ribosomal subunits in an open conformation, 95673 particles with a 40S small ribosomal subunit in a half-closed conformation and 76966 particles with 48S PICs in a closed head conformation (**Extended Data Fig. 1**). The particles with PICs were further classified focused on the ternary complex resulting in 25852 particles without tRNA_i_ or eIF2 and 51031 particles with tRNA_i_ and eIF2 (**Extended Data Fig. 1**). The particles with tRNA_i_ and eIF2 were re-extracted with and coarsened to 1.44 Å/pixels and the structure refined to 3.1 Å resolution (**Extended Data Fig. 1-3**). This particle pool was also further classified for finer 40S head movements which identified a more populated state of 48S complexes containing 39810 particles that were used for further analysis and a less populated state containing 10877 particles. The particle pool with 39810 particles was refined to a map at 2.9 Å resolution (**Extended Data Fig. 1-3**) followed by multibody refinement, which resulted in a map for the 40S body including mRNA and eIF1A at a resolution of 3.0 Å, a map for the 40S head including the ternary complex at a resolution of 3.0 Å and a map containing eIF3 and ABCE1 at a resolution of 6.5 Å (**Extended Data Fig. 1**). The maps containing 40S body and 40S head were further improved by signal subtraction to obtain better map definition and slightly better resolution of 2.9 Å and 2.9 Å (**Extended Data Figs. 1-3**). These two maps derived from signal subtraction with centering were amplitude-corrected using Phenix auto_sharpen, combined to a composite map using the software CHIMERA, and used for map interpretation and atomic model building.

After 2D classification 72549 good particles of the Kozak 48S PICs (ACCAUGG mRNA) containing the eIF1A W70A mutant were extracted and coarsened to 4.32 Å/pixels. Focused classification on the head of the 40S small ribosomal subunit resulted in 42682 particles containing the 40S small ribosomal subunits in an open conformation, 8457 particles with a 40S small ribosomal subunit in a half-closed conformation and 55283 particles with 48S PICs and a closed head conformation (**Extended Data Fig. 1**). The particles with PICs were further classified focused on the ternary complex resulting in 24556 particles without tRNA_i_ or eIF2 and 30727 particles with tRNA_i_ and eIF2 (**Extended Data Fig. 1**). The particles with tRNA_i_ and eIF2 were re-extracted with at the full sampling of 0.72 Å/pixels and refined to 3.2 Å resolution (**Extended Data Fig. 1-3**). Multibody refinement resulted in a map for the 40S body including mRNA and eIF1A at a resolution of 3.2 Å, a map for the 40S head including the ternary complex at a resolution of 3.3 Å and a map containing eIF3 and ABCE1 at a resolution of 7.0 Å (**Extended Data Figs. 1 & 2**). This complex was not classified for fine head movements because the mRNA and eIF1A side-chains were well visible without further classification. Further sorting confirmed that the mRNA does not form different conformations in the A-site. Therefore, the 40S body and head maps and masks were directly used further for signal subtraction and centering followed by focused refinement, which resulted in improved maps at 3.2 Å for both regions (**Extended Data Figs. 1-3**). These maps were amplitude-corrected using Phenix auto_sharpen^66^, combined to a composite map using CHIMERA, and used for atomic model building and map interpretations. Resolution estimation by Fourier shell correlation (FSC^67–69)^ calculations using the 0.143 criterion were done with the software Relion^63^ and map-model correlation calculations were done with the software Phenix^66^. All FSC curves were plotted using Python. Orientational distribution plots were prepared using the programme VUE^70^.

### Model fitting and refinement

The high-quality maps enabled building and refining atomic models with good geometrical parameters (**Extended Data Table 1**). For this, the high-resolution atomic model for the 40S ribosomal subunit of the human 80S ribosome containing rRNA modifications (PDB ID: 8QOI^34^) was fitted into the cryo-EM-map using the software Coot^71^. Atomic models for the tRNA_i_ were derived from tRNA_i_ coordinates of PDB ID 3V11^72^. The chemical modifications of *Homo sapiens* tRNA_i_ were added according to the MODOMICS database^73,74^. The atomic model of the ternary complex was generated by manually adapting alpha fold models for eIF2, where the structure of the archaea ternary complex^72^ was used as a guidance in less well resolved parts of the map, such as the acceptor arm of the tRNA, domain 3 of eIF2α (according to the Uniprot database^75^ the numbering of sequence stretch of eIF2α is Arg53, Arg54, Arg55 and Arg57, with Arg55 being the only residue interacting with the mRNA) as well as eIF2β and eIF2γ. Atomic models of eIF3 and eIF1A were generated using the Alphafold option of the Phenix software “build and refine” and adjusted with the software Coot. The atomic model was refined iteratively using the software Coot and Phenix^76–78^ until a good model geometry was obtained (**Extended Data Table 2**). Map interpretation and fitting took into account the role of charges in the cryo-EM density maps which are electrostatic potential maps: positively charged atoms tend to be better resolved than negatively charged atoms^79–81;^ for example the series of arginine residues of eIF2α next to mRNA, 18S rRNA and tRNA_i_. Figures were prepared using the software ChimeraX^82^.

The tRNA_i_ shows classical Watson-Crick base pairs that form the codon-anticodon mini helix (**Fig. 1D**), complemented by characteristic direct interactions of the tRNA and with the N6 and N7 position of the A+1 nucleotide base of the mRNA via the carboxy moiety of t_6_A37. Additionally, the hydroxyl group of t_6_A forms hydrogen bonds with the 2’OH moiety of C-1 base of the mRNA, thereby further stabilizing the conformation of the mRNA-tRNA duplex. Additional interactions of the mRNA backbone (nucleotides - 1 to +4) with the 18S rRNA contribute to stabilize the mRNA in the complex. The codon-anticodon mini-helix is kept in place by two key 18S rRNA residues, C1701 and 1-methyl-3-(3-amino-3-carboxylpropyl)Ψ1248 (m^1^acp^3^Ψ1248)^21,39^, which are bridged together through hydrogen bonds and contact the C34 (tRNA) backbone (**Extended Data Fig. 5F**).

