## Extended Figures for "Translation initiation by the Kozak mRNA sequence is based on a conformational readout on the ribosome"

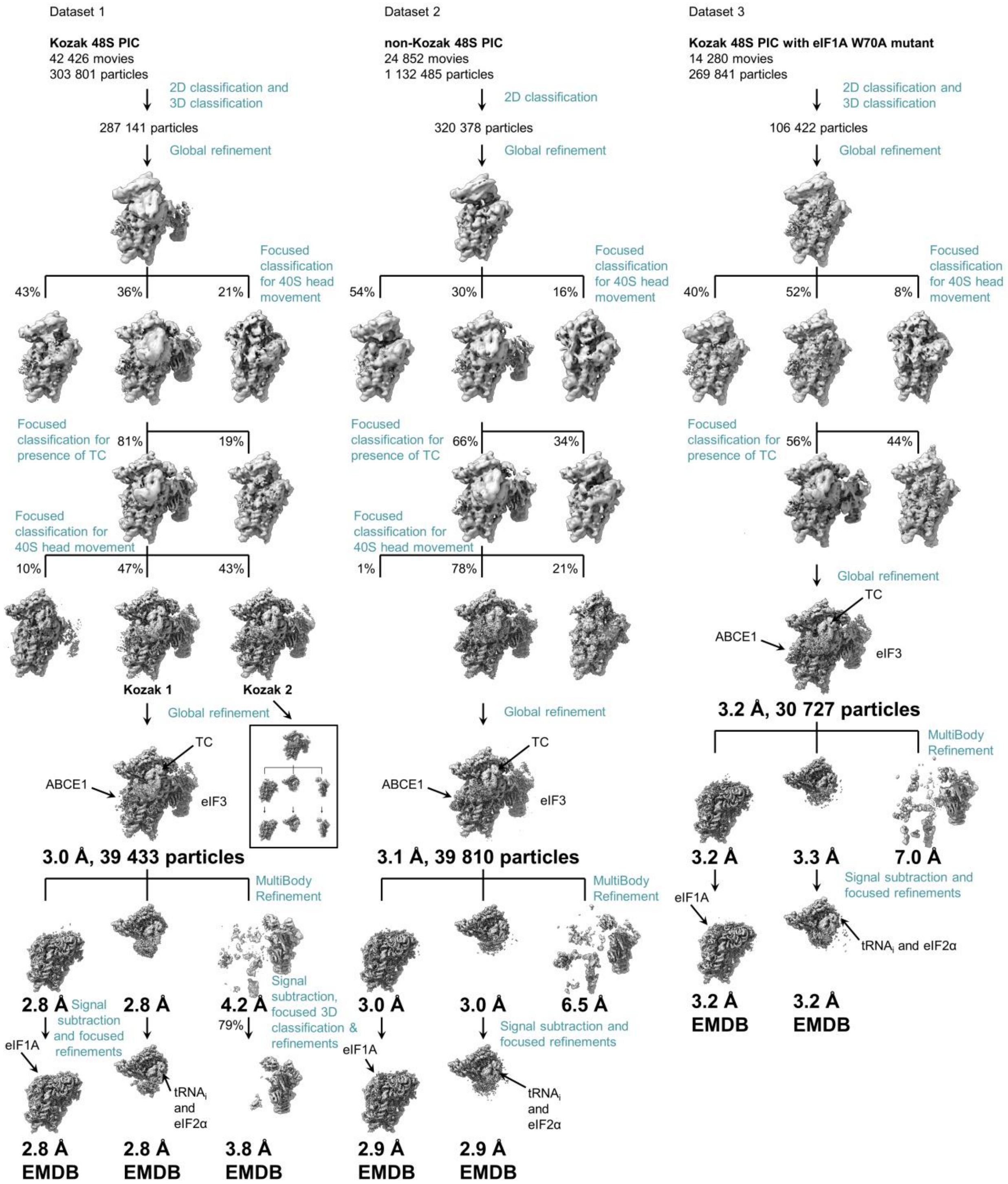
Dataset 1
Kozak 48S PIC
42 426 movies
303 801 particles
2D classification and 3D classification
287 141 particles
Global refinement
Focused classification for 40S head movement
43%
36%
21%
Focused classification for presence of TC
81%
19%
Focused classification for 40S head movement
10%
47%
43%
Kozak 1
Kozak 2
Global refinement
TC
ABCE1
eIF3
3.0 Å, 39 433 particles
MultiBody Refinement
2.8 Å
Signal subtraction and focused refinements
2.8 Å
4.2 Å
Signal subtraction, focused 3D classification & refinements
79%
eIF1A
tRNAi and eIF2α
2.8 Å EMDB
2.8 Å EMDB
3.8 Å EMDB
Dataset 2
non-Kozak 48S PIC
24 852 movies
1 132 485 particles
2D classification
320 378 particles
Global refinement
Focused classification for 40S head movement
54%
30%
16%
Focused classification for presence of TC
66%
34%
Focused classification for 40S head movement
1%
78%
21%
Global refinement
TC
ABCE1
eIF3
3.1 Å, 39 810 particles
MultiBody Refinement
3.0 Å
3.0 Å
6.5 Å
Signal subtraction and focused refinements
eIF1A
tRNAi and eIF2α
2.9 Å EMDB
2.9 Å EMDB
Dataset 3
Kozak 48S PIC with eIF1A W70A mutant
14 280 movies
269 841 particles
2D classification and 3D classification
106 422 particles
Global refinement
Focused classification for 40S head movement
40%
52%
8%
Focused classification for presence of TC
56%
44%
Global refinement
TC
ABCE1
eIF3
3.2 Å, 30 727 particles
MultiBody Refinement
3.2 Å
3.3 Å
7.0 Å
Signal subtraction and focused refinements
eIF1A
tRNAi and eIF2α
3.2 Å EMDB
3.2 Å EMDB

**Sorting scheme of the cryo-EM data for the Kozak 48S PIC (dataset 1, left), the non-Kozak 48S PIC (dataset 2, middle) and the Kozak 48S PIC with the eIF1A W70A mutant (dataset 3, right)**

The data were first sorted for major conformational changes in the 40S head with respect to the 40S body. The particles in the “closed” head conformation were then further sorted for TC occupancy. A second classification for fine differences in the head conformations was done for dataset 1 and 2 but not for dataset 3 which appeared to have only one mRNA conformation. The refined maps were then subjected to multibody refinement followed by signal subtraction and focused refinement on the individual bodies.

**Extended Data Fig. 1**

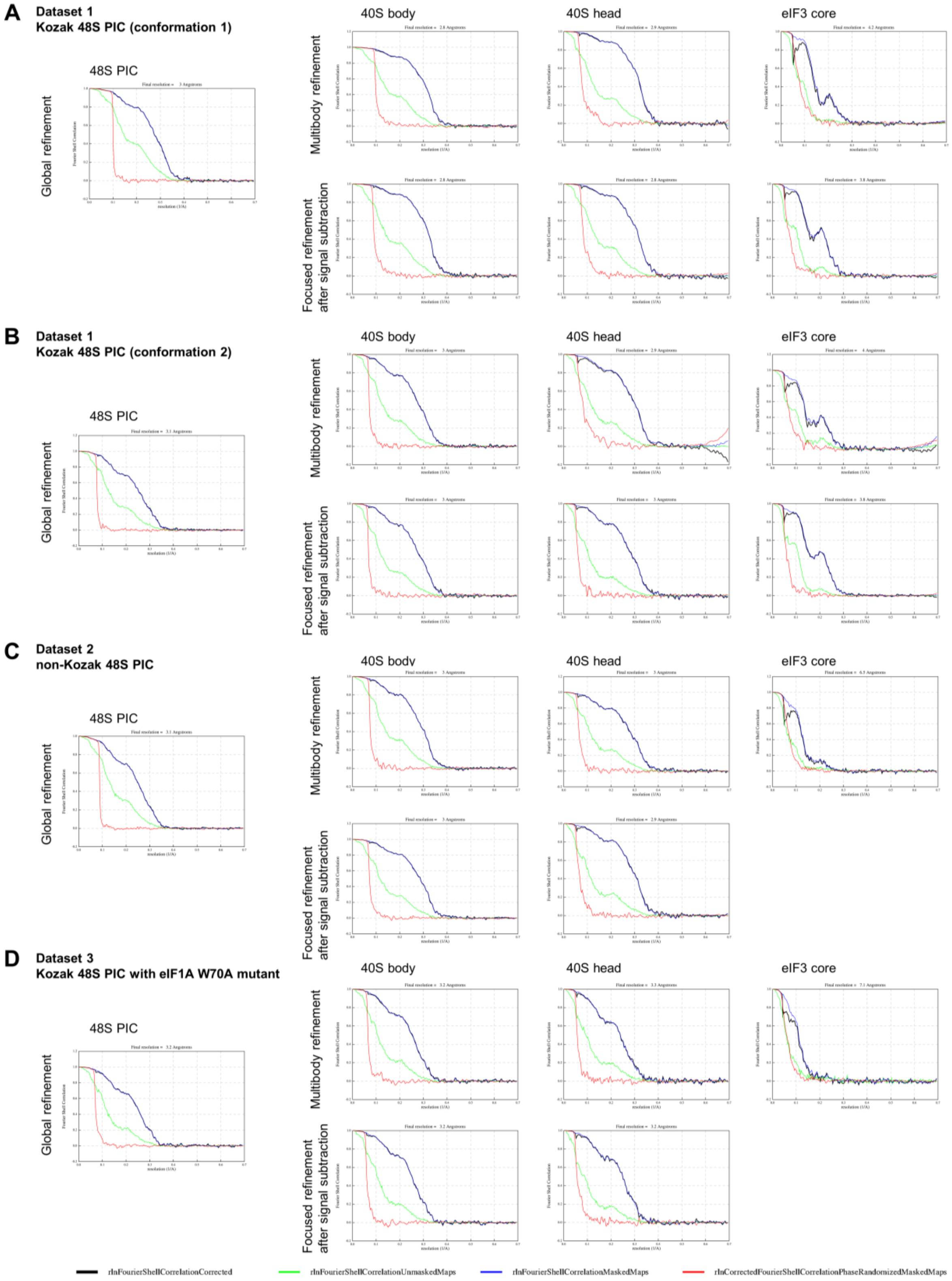


**FSC curves for the various cryo-EM 3D reconstructions**

**(A-D)** FSC curves for the various global, focused and multi-body refinements of datasets 1-3.

**Extended Data Fig. 2**

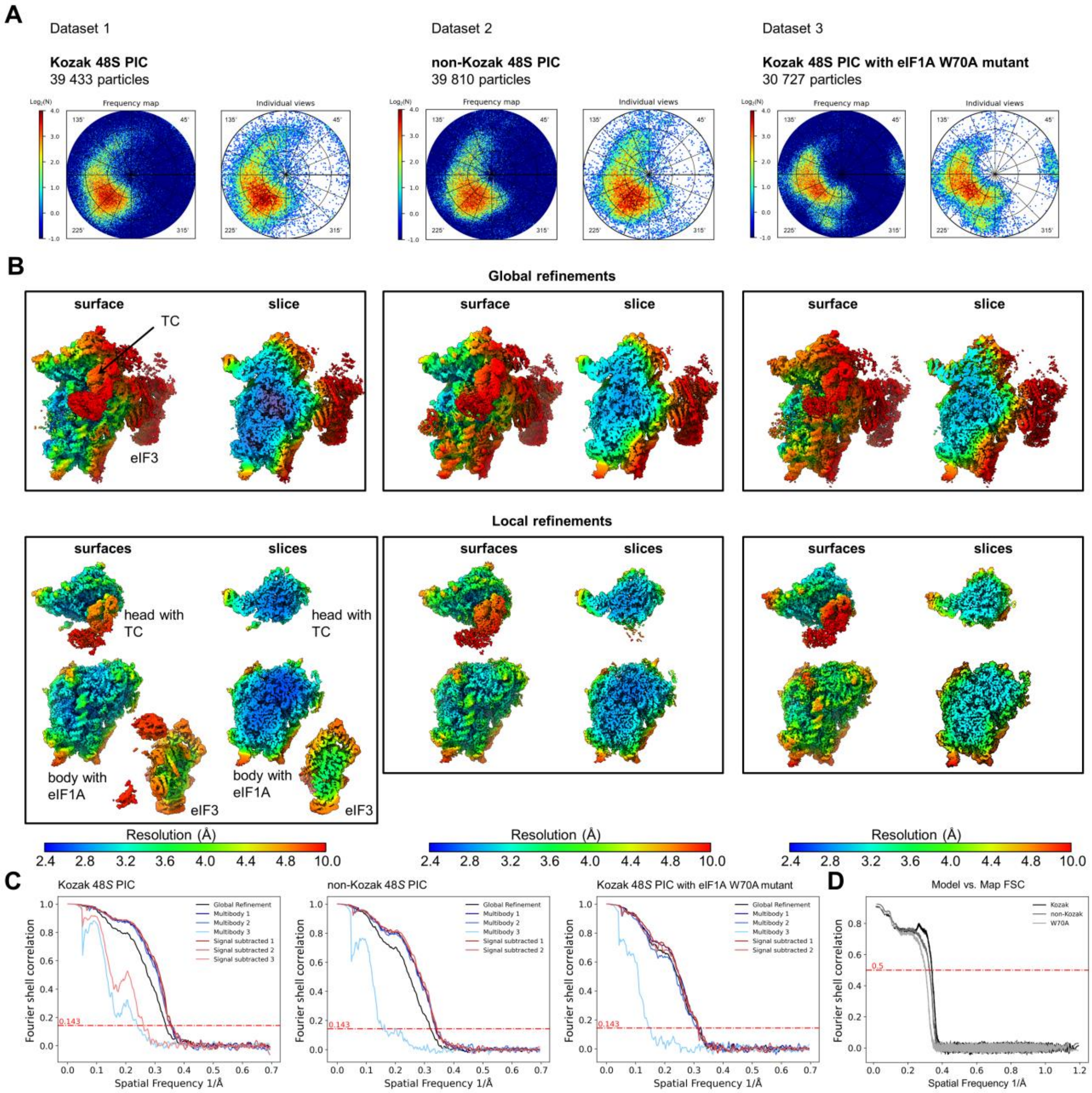
A
Dataset 1
Kozak 48S PIC
39 433 particles
Dataset 2
non-Kozak 48S PIC
39 810 particles
Dataset 3
Kozak 48S PIC with eIF1A W70A mutant
30 727 particles
Frequency map
Individual views
B
Global refinements
surface
slice
TC
eIF3
Local refinements
surfaces
slices
head with TC
body with eIF1A
eIF3
Resolution (Å)
2.4 2.8 3.2 3.6 4.0 4.4 4.8 10.0
C
Kozak 48S PIC
non-Kozak 48S PIC
Kozak 48S PIC with eIF1A W70A mutant
Global Refinement
Multibody 1
Multibody 2
Multibody 3
Signal subtracted 1
Signal subtracted 2
Signal subtracted 3
Fourier shell correlation
Spatial Frequency 1/Å
0.143
D
Model vs. Map FSC
Kozak
non-Kozak
W70A
0.5

**Orientational distribution and resolution of cryo-EM reconstructions**

**(A)** Orientational distribution of particles for the Kozak 48S PIC (left), the non-Kozak 48S PIC (middle) and the Kozak 48S PIC with the eIF1A W70A mutant (right) as calculated with the *VUE* program using the Lambert projection[70]. For the plots the hemisphere was split into 3600 cells of an equal surface area corresponding to division of the spherical half-meridian in 30 steps, by 3° each. Coverage of the hemisphere is 78% for dataset 1, 76% for dataset 2 and 69% for dataset 3.

**(B)** Local resolution distribution for the globally refined maps and the focused refined maps after signal subtraction. The scale bar is coloured according to the respective local resolution (Å).

**(C)** FSC curves for all three datasets for globally refined maps, maps after multibody refinement and focused refined maps after signal subtraction. The curves were calculated using the Relion software and plotted using Python. According to the 0.143 cutoff criterium the estimated resolutions are for the Kozak 48S PIC 3.0 Å for the globally refined map, 2.8 Å after multibody refinement of the 40S body (multibody 1), 2.8 Å after multibody refinement of the 40S head (multibody 2) and 4.2 Å after multibody refinement of eIF3 (multibody 3). The map quality was further improved by signal subtraction and focused refinement reaching resolution values of 2.8 Å for the 40S body (signal subtracted 1), 2.8 Å for the 40S head (signal subtracted 2) and 3.9 Å after multibody refinement of eIF3 (signal subtracted 3).

Corresponding estimated resolution values for the non-Kozak 48S PIC are: 3.1 Å for the globally refined map, 3.0 Å after multibody refinement of the 40S body (multibody 1), 3.0 Å after multibody refinement of the 40S head (multibody 2) and 6.5 Å after multibody refinement of eIF3 (multibody 3). The map qualities of the 40S head and body were further improved by signal subtraction and focused refinement reaching resolution values of 2.9 Å for the 40S body (signal subtracted 1) and 2.9 Å for the 40S head (signal subtracted 2).

Corresponding estimated resolution values for the Kozak 48S PIC with the eIF1A W70A mutant are: 3.2 Å for the globally refined map, 3.2 Å after multibody refinement of the 40S body (multibody 1), 3.3 Å after multibody refinement of the 40S head (multibody 2) and 7.0 Å after multibody refinement of eIF3 (multibody 3). The map qualities of the 40S head and body were further improved by signal subtraction and focused refinement reaching resolution values of 3.2 Å for the 40S body (signal subtracted 1) and 3.2 Å for the 40S head (signal subtracted 2). See Table 1 for more details.

**(D)** Model vs. map FSC curves for the atomic models built into the composite maps derived from focused refinements after signal subtraction for the Kozak 48S PIC (Kozak), the non-Kozak 48S PIC (non-Kozak) and the Kozak 48S PIC with the eIF1A W70A mutant (W70A); reported resolution values using a 0.5 cutoff are 3.1 Å for the Kozak 48S PIC, 3.2 Å for the non-Kozak 48S PIC and 3.7 Å for the Kozak 48S PIC with the eIF1A W70A mutant. See Table 1 for more details.

**Extended Data Fig. 3**

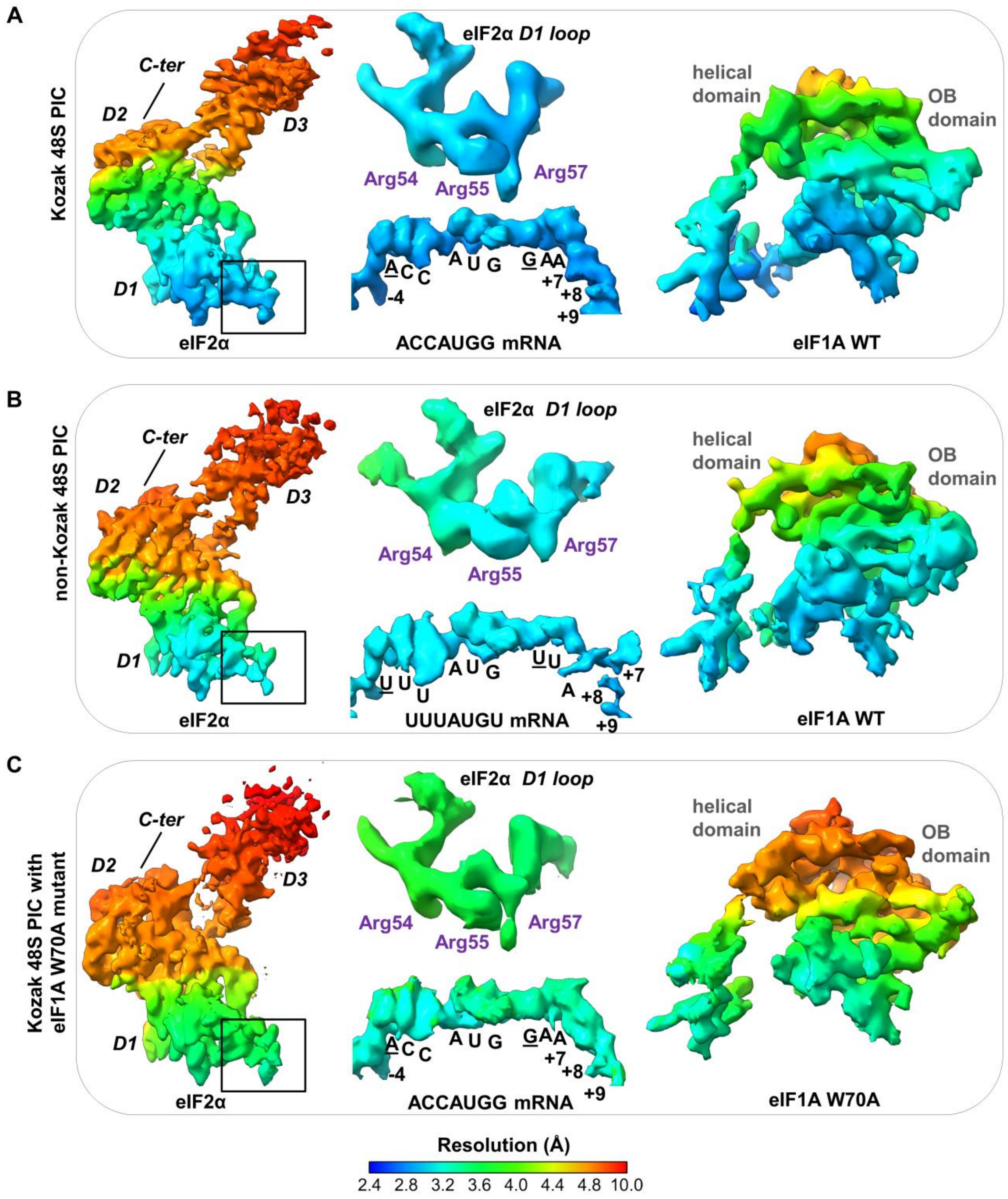
A
Kozak 48S PIC
C-ter
D2
D3
D1
eIF2α
eIF2α D1 loop
Arg54
Arg55
Arg57
ACCAUGG mRNA
helical domain
OB domain
eIF1A WT
B
non-Kozak 48S PIC
UUUAUGU mRNA
eIF1A WT
C
Kozak 48S PIC with eIF1A W70A mutant
eIF1A W70A
Resolution (Å)
2.4 2.8 3.2 3.6 4.0 4.4 4.8 10.0

**Local resolution distribution in the eIF2α, mRNA and eIF1A parts of the complexes**

**(A)** Surface colour representation of local resolution distribution in the Kozak 48S PIC with focus on eIF2α (left), D1 loop of eIF2α and mRNA (middle) and eIF1A (right).

**(B)** Surface colour representation of local resolution distribution in the non-Kozak 48S PIC with focus on eIF2α, D1 loop of eIF2α, mRNA and eIF1A as in (A).

**(C)** Surface colour representation of local resolution distribution in the Kozak 48S PIC with mutant eIF1A W70A as in (A&B).

**Extended Data Fig. 4**

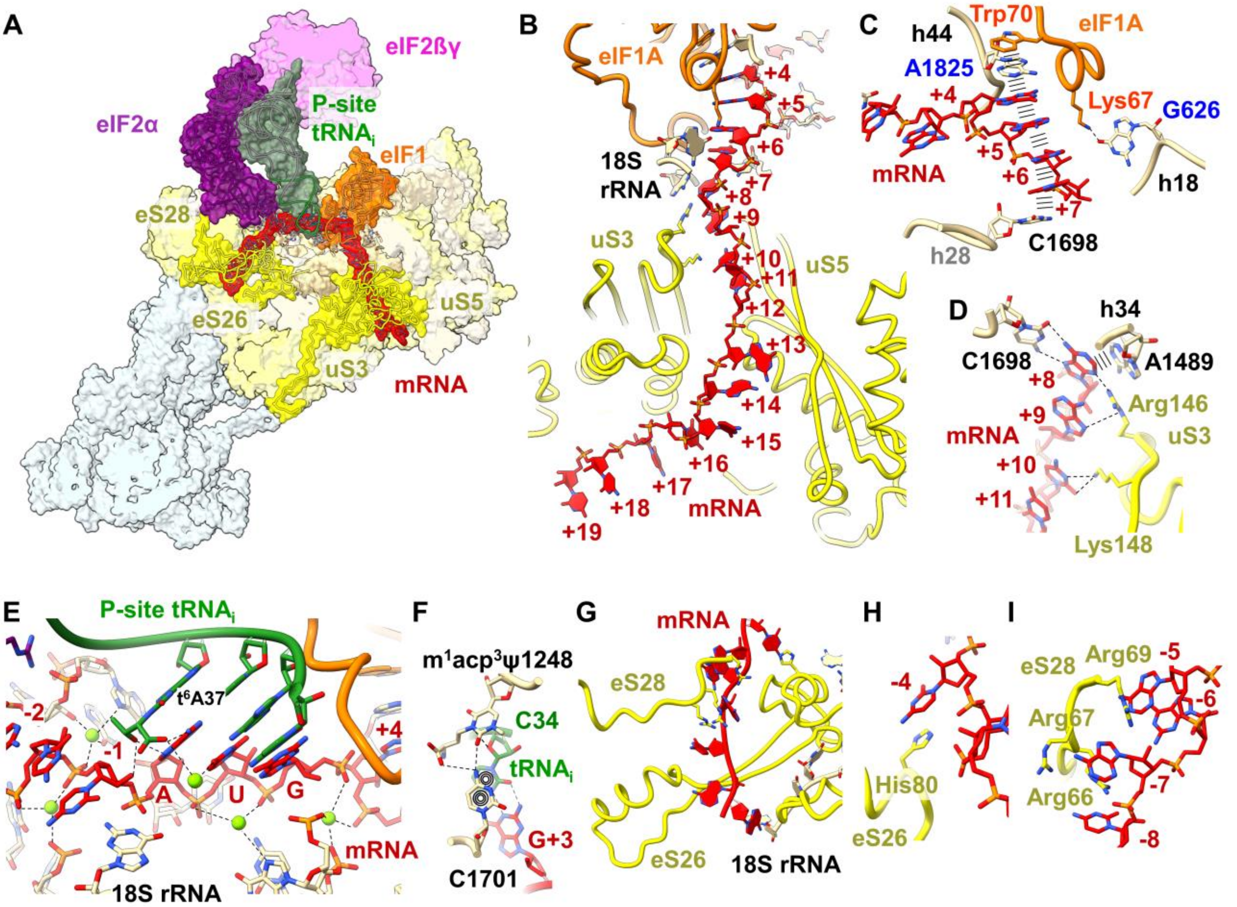
A
eIF2ßγ
eIF2α
P-site
tRNA$_i$
eIF1
eS28
eS26
uS5
uS3
mRNA
B
eIF1A
+4
+5
+6
18S
rRNA
+7
+8
+9
uS3
+10
+11
uS5
+12
+13
+14
+15
+16
+17
+18
mRNA
+19
C
h44
Trp70
eIF1A
A1825
+4
Lys67
G626
mRNA
+5
+6
h18
+7
h28
C1698
D
h34
C1698
A1489
+8
+9
Arg146
mRNA
uS3
+10
+11
Lys148
E
P-site tRNA$_i$
t$^6$A37
-2
-1
A
U
G
+4
mRNA
18S rRNA
F
m$^1$acp$^3$ψ1248
C34
tRNA$_i$
G+3
C1701
G
mRNA
eS28
eS26
18S rRNA
H
-4
His80
eS26
I
eS28
Arg69
-5
-6
Arg67
Arg66
-7
-8

**Interactions of the mRNA with rRNA, tRNA$_i$ and initiation factors eIF1A and eIF2α**

**(A)** Cryo-EM structure of the Kozak 48S PIC at 2.8 Å resolution as seen from the front, sliced to focus on the mRNA binding tunnel in the 40S small ribosomal subunit. Ribosomal proteins that interact with the mRNA are highlighted.

**(B)** Overview over the mRNA strand in the mRNA entry tunnel with uS3, uS5 and 18S rRNA as main interacting partners.

**(C)** Zoom on mRNA bases +4 to +7 which stack with each other. Further stabilization occurs via π-stacking with A1825 which is flipped out and inserts between Trp70 and G+4. C1698 of the 18S rRNA stabilizes the mRNA via π-stacking with C+7 of the mRNA. Lys67 is part of the α1-helix of eIF1A and interacts with decoding residue G626.

**(D)** Zoom on mRNA bases +8 to +11 which are positioned by π-stacking with A1489 and non-Watson-Crick base-pairing with C1698 of the 18S rRNA. Further stabilization is provided by Arg146 uS3, which interacts with A+8 and A+9 as well as Lys148 of ribosomal protein uS3, which interacts with C+10.

**(E)** Zoom on the mRNA binding site at the E-, P- and A-site of the 40S small ribosomal subunit in the human 48S PIC. The AUG start codon base-pairs with the tRNA$_i$ anti-codon loop. The positioning of the RNA phosphate moieties of the mRNA are stabilized via ions that bridge to various nucleotide bases of the 18S rRNA as well as to the carboxyl group of the threonylcarbamyl modification of A37 of the P-site tRNA$_i$. The keto group of the threonylcarbamyl modification of A37 is interacting with the 2'OH moiety of mRNA base C-1 as well as with N7 of the mRNA A +1 giving further stabilization for the AUG start codon recognition by tRNA$_i$.

**(F)** Interactions of $m^1acp^3\Psi$1248 with C1701 and the G+3 / C34 mRNA codon P-site tRNA anticodon pair.

**(G)** Overview over the mRNA bound to the mRNA exit tunnel with eS26, eS28 and the 3' end of the 18S rRNA as main interacting partners.

**(H)** Interaction of C-4 with His80 of eS26 via π-stacking.

**(I)** Interaction of A-5 to A-7 with eS28 via π-stacking with residue Arg69 and hydrogen bonds of Arg66 with the mRNA phosphate backbone.

**Extended Data Fig. 5**

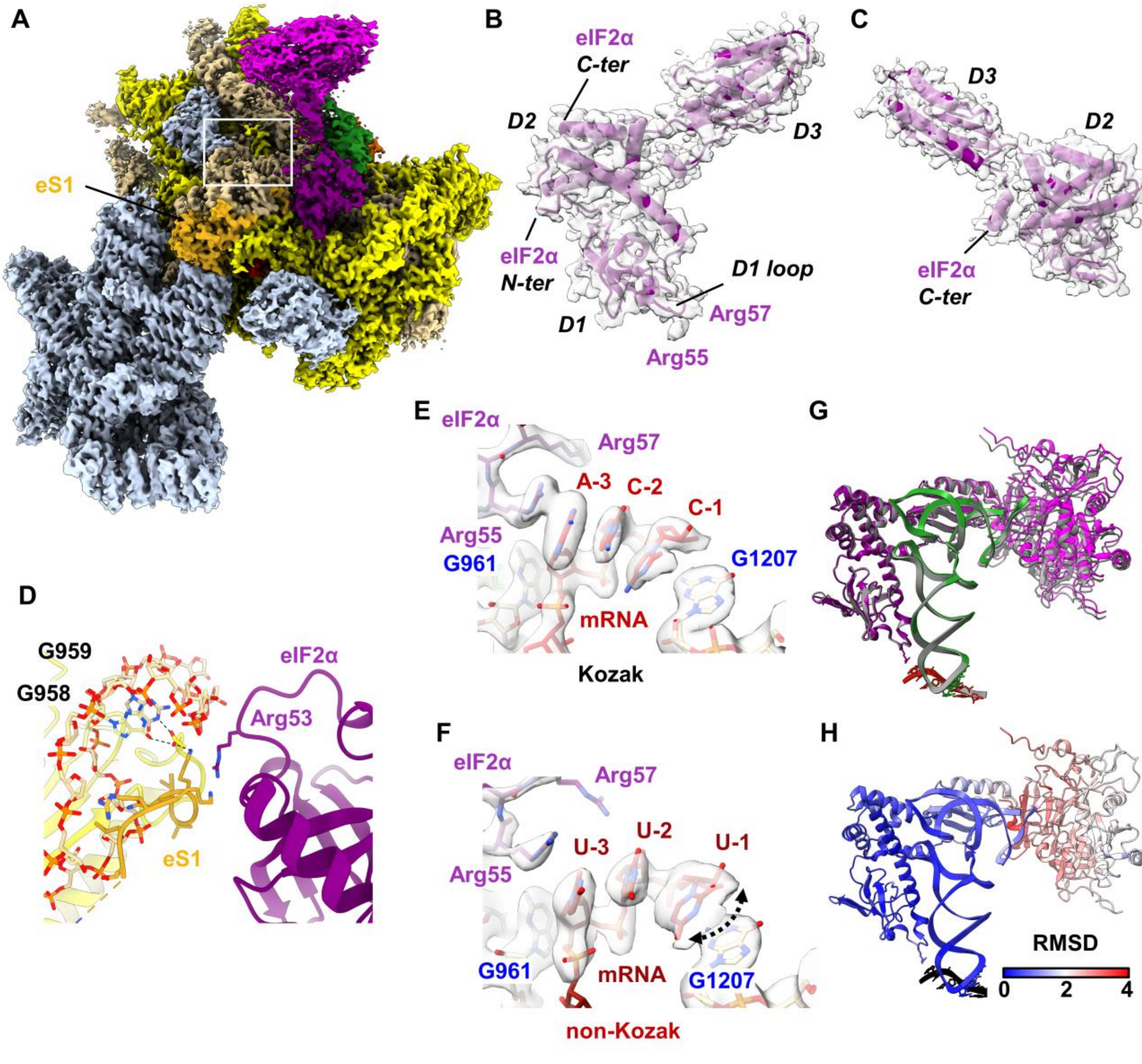
A
eS1
B
eIF2α
C-ter
D2
D3
eIF2α
N-ter
D1 loop
D1
Arg57
Arg55
C
D3
D2
eIF2α
C-ter
D
G959
G958
eIF2α
Arg53
eS1
E
eIF2α
Arg57
A-3
C-2
C-1
Arg55
G961
G1207
mRNA
Kozak
F
eIF2α
Arg57
U-3
U-2
U-1
Arg55
G961
mRNA
G1207
non-Kozak
G
H
RMSD
0
2
4

**Details of eIF2α and its binding within the human 48S PICs**

**(A)** Overview of the human 48S PIC. eS1 is highlighted in orange. The white box marks the place where the eS1 N-terminus is bound between eIF2α (purple), uS11 (yellow) and the 18S rRNA (light beige).

**(B,C)** Overview of eIF2α in the structure showing the clearly visible C-terminal α-helix that folds back onto domain 2 (D2).

**(D)** Zoomed view on the eS1 N-terminus (orange).

**(E)** Surface representation of the upstream part of the mRNA sequence recognition by a preformed G-clamp and eIF2α in the Kozak 48S PIC as shown in **Fig. 2D**.

**(F)** Surface representation of the upstream part of the mRNA sequence recognition by a preformed G-clamp in the non-Kozak 48S PIC as shown in **Fig. 2E.** Broader density of the C-1 base is indicated by an arrow.

**(G)** overlay of the TC in the Kozak 48S PIC (colored: $tRNA_i$: green, eIF2α: purple, mRNA: black, eIF2βγ: magenta) and the non-Kozak 48S PIC (gray).

**(H)** RMSD (root mean square deviation) calculation of the difference between the TC in the Kozak 48S PIC and the TC in the non-Kozak 48S PIC.

**Extended Data Fig. 6**

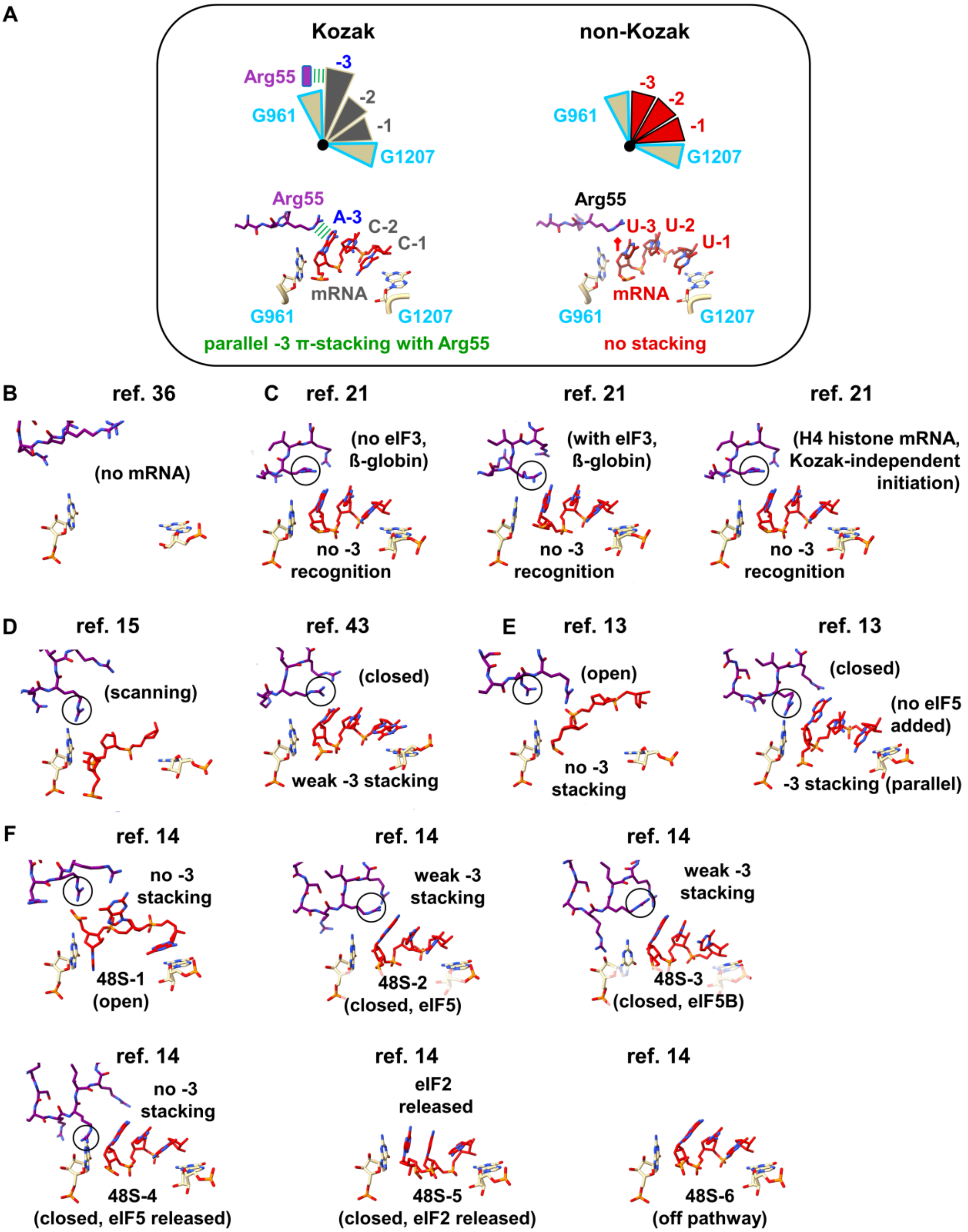
A
Kozak
non-Kozak
Arg55
-3
-2
-1
G961
G1207
A-3
C-2
C-1
mRNA
parallel -3 π-stacking with Arg55
U-3
U-2
U-1
no stacking
B
ref. 36
(no mRNA)
C
ref. 21
(no eIF3, ß-globin)
no -3 recognition
(with eIF3, ß-globin)
(H4 histone mRNA, Kozak-independent initiation)
D
ref. 15
(scanning)
ref. 43
(closed)
weak -3 stacking
E
ref. 13
(open)
no -3 stacking
(closed)
(no eIF5 added)
-3 stacking (parallel)
F
ref. 14
48S-1 (open)
48S-2 (closed, eIF5)
48S-3 (closed, eIF5B)
48S-4 (closed, eIF5 released)
eIF2 released
48S-5 (closed, eIF2 released)
48S-6 (off pathway)

**Comparisons of eukaryotic 48S PICs (-3 region)**

Shown is the same close-up view onto the -3 nucleotide region of the mRNA as in Figure 2D&E to compare the positioning of Arg55 (eIF2α) observed in this study with previously published structures of mammalian 48S PICs.

**(A)** Interactions of eIF2α and the mRNA as observed in this study for Kozak 48S PIC and non-Kozak 48S PIC, revealing strong π-stacking (parallel configuration of the guanidinium moiety and the aromatic purine) versus no interaction with a pyrimidine, respectively.

**(B)** Close-up view on the -3 region of the human 43S PIC (PDB: 7A09).

**(C)** Close-up view on the -3 region of the rabbit 48S PIC with either the Kozak-dependent beta globin ((PDB: 6YAL, 6YAM) mRNA or the Kozak-independent histone H4 mRNA (PDB: 6YAN).

**(D)** Close-up view on the -3 region of the scanning human 48S PIC (PDB: 6ZMW) and in the closed state (PDB: 8OZ0).

**(E)** Close-up view on the -3 region of the human 48S PIC in open (PDB: 7QP6) and in the closed state (PDB: 7QP7).

**(F)** Close-up view on the -3 region of the human 48S PIC with eIF1 in open (PDB: 8PJ1) and in closed states (PDB: 8PJ2, 8PJ3, 8PJ4, 8PJ5, 8PJ6).

**Extended Data Fig. 7**

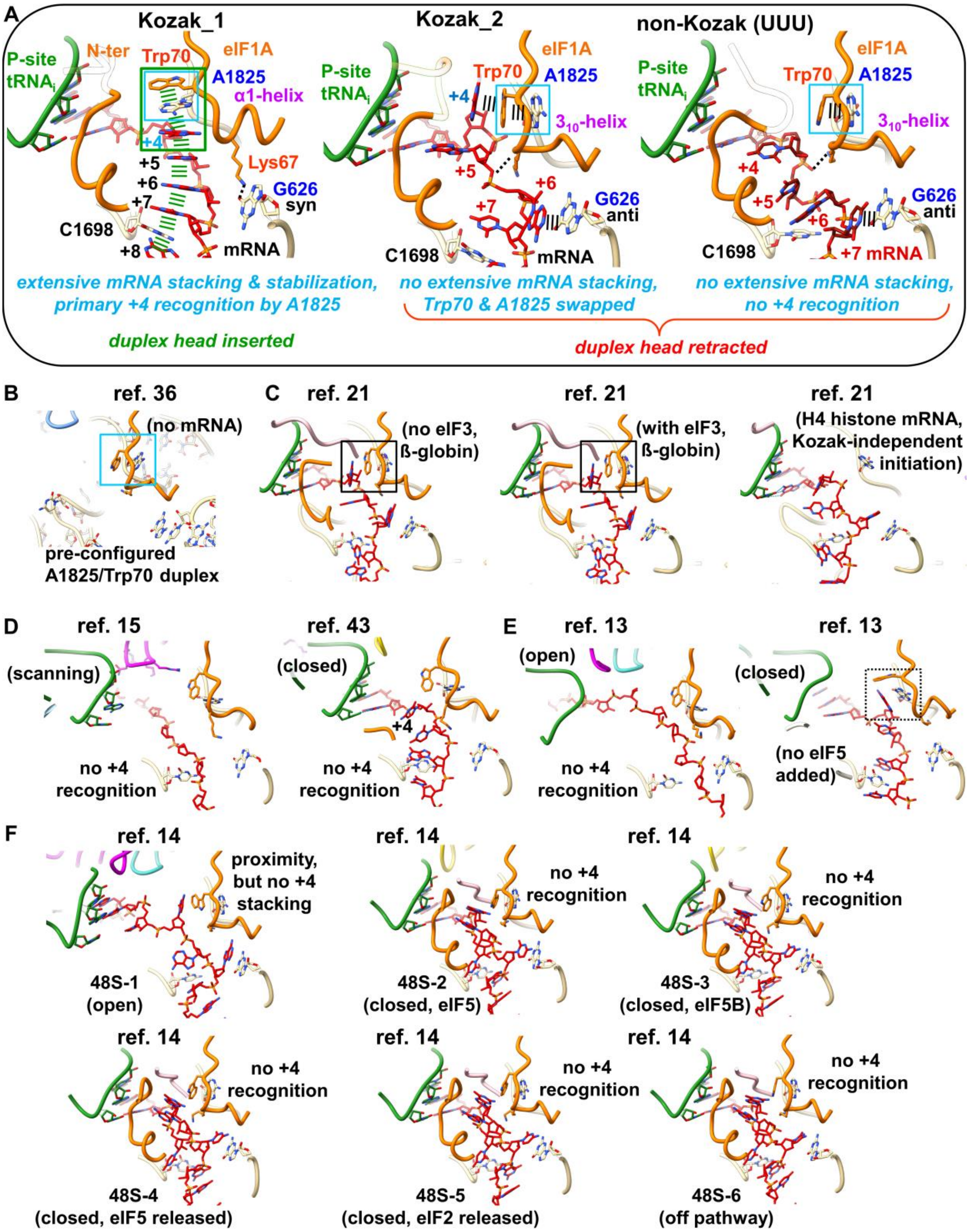
A
Kozak_1
Kozak_2
non-Kozak (UUU)
P-site tRNAi
N-ter
Trp70
eIF1A
A1825
α1-helix
3₁₀-helix
+4
+5
+6
+7
+8
Lys67
G626
syn
anti
C1698
mRNA
extensive mRNA stacking & stabilization, primary +4 recognition by A1825
no extensive mRNA stacking, Trp70 & A1825 swapped
no extensive mRNA stacking, no +4 recognition
duplex head inserted
duplex head retracted
B
ref. 36
(no mRNA)
pre-configured A1825/Trp70 duplex
C
ref. 21
(no eIF3, ß-globin)
(with eIF3, ß-globin)
(H4 histone mRNA, Kozak-independent initiation)
D
ref. 15
(scanning)
no +4 recognition
ref. 43
(closed)
E
ref. 13
(open)
(no eIF5 added)
F
ref. 14
proximity, but no +4 stacking
48S-1 (open)
48S-2 (closed, eIF5)
48S-3 (closed, eIF5B)
48S-4 (closed, eIF5 released)
48S-5 (closed, eIF2 released)
48S-6 (off pathway)

### Comparisons of eukaryotic 48S PICs (+4 region)

Shown is the same close-up view onto the A-site as in Figure 3A to compare the Trp70/A1825 duplex conformations observed in this study with previously published structures of mammalian 48S PICs.

**(A)** Interactions of eIF1A and the mRNA as observed in this study for Kozak 48S PIC and non-Kozak 48S PIC.

**(B)** Close-up view on the A-site of the human 43S PIC (PDB: 7A09).

**(C)** Close-up view on the A-site of the rabbit 48S PIC with either the Kozak-dependent beta globin ((PDB: 6YAL, 6YAM) mRNA or the Kozak-independent histone H4 mRNA (PDB: 6YAN).

**(D)** Close-up view on the A-site of the scanning human 48S PIC (PDB: 6ZMW) and in the closed state (PDB: 8OZ0).

**(E)** Close-up view on the A-site of the human 48S PIC in open (PDB: 7QP6) and in the closed state (PDB: 7QP7).

**(F)** Close-up view on the A-site of the human 48S PIC with eIF1 in open (PDB: 8PJ1) and in closed states (PDB: 8PJ2, 8PJ3, 8PJ4, 8PJ5, 8PJ6).

### Extended Data Fig. 8

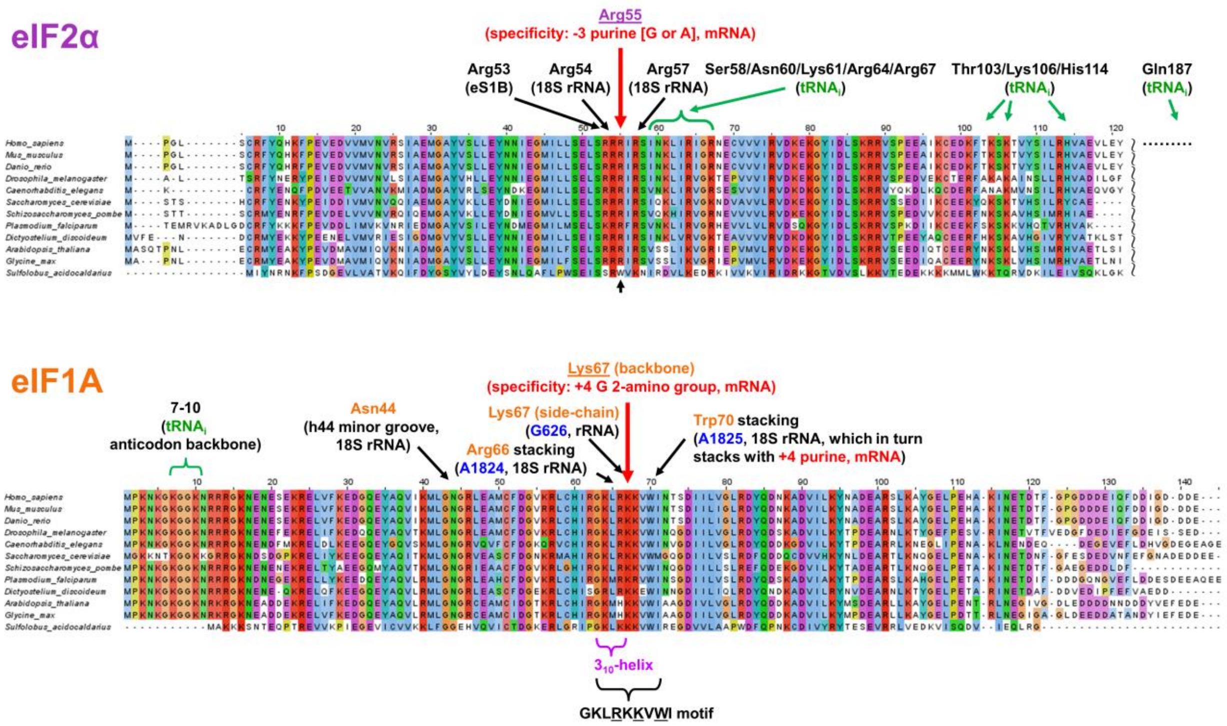


**Sequence conservation of eIF1A and eIF2α**

Multiple sequence alignments of eIF2α (top) and eIF1A (bottom). Key interacting residues in the Kozak 48S PIC are annotated.

**Extended Data Fig. 9**

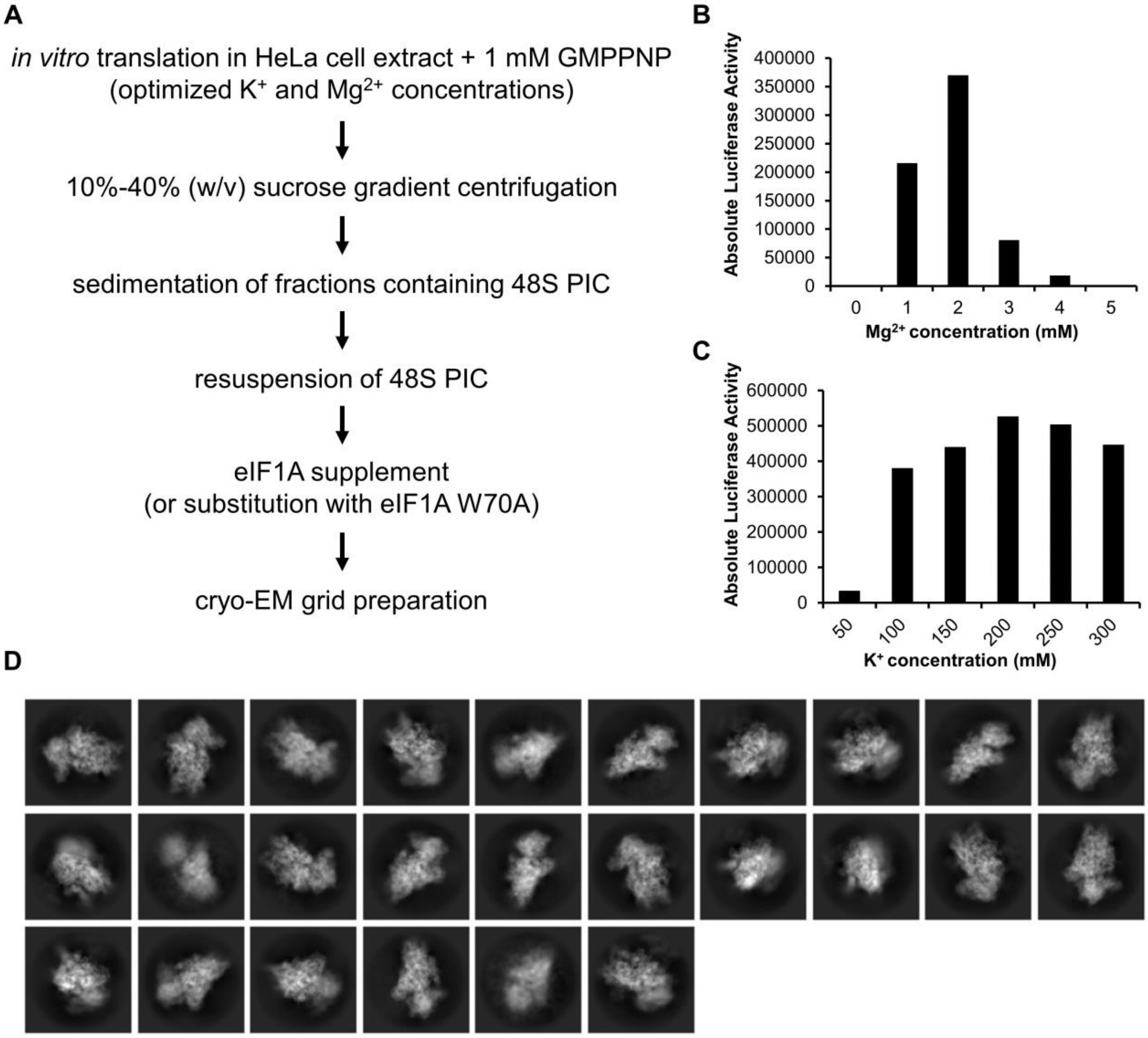


**Sample purification of 48S PICs**

**(A)** Flow scheme of 48S PIC sample purification.

**(B)** Optimization of $Mg^{2+}$ (magnesium acetate) salt concentrations with constant concentration of 120 mM $K^{+}$. Shown are average values from absolute luciferase readout values of two technical duplicates.

**(C)** Optimization of $K^{+}$ (potassium acetate) salt concentrations with constant concentration of 2 mM $Mg^{2+}$. Shown are average values from absolute luciferase readout values of two technical duplicates.

**(D)** Representative class averages of 48S PIC particles.

**Extended Data Fig. 10**

| **Image processing** | **Data set 1** | **Data set 2** | **Data set 3** |
| --- | --- | --- | --- |
| Total number of extracted particles | 303 801 | 1 132 485 | 106 422 |
| Final number of particles used | 39 433 | 39 810 | 30 727 |
| Pixel size (Å) | 0.72 | 0.72 | 0.72 |
| Box size (pixels) | 480 | 480 | 480 |
| Symmetry | C1 | C1 | C1 |
| Map sharpening B-factor (Å$^2$) | -39.3 (40S body+eIF1A+mRNA) -39.2 (40S head +TC) -124.7 (eIF3) | -30.0 (40S body+eIF1A+mRNA) -31.7 (40S head +TC) | -45.7 (40S body+eIF1A+mRNA) -50.7 (40S head +TC) |
| Global map resolution (Å) | 2.8; 2.8; 3.8 | 2.9; 2.9 | 3.2; 3.2 |

**Table 1 Cryo-EM data collection and structure determination**

| Atomic model refinement | Data set 1 | Data set 2 | Data set 3 |
|---|---|---|---|
| Model composition<br>Chains<br>Atoms<br>Residues (a.a. / nucleotides)<br>Ions | <br>56<br>108 849<br>9 304 / 1841<br>112 | <br>45<br>85 799<br>5 795 / 1 841<br>92 | <br>45<br>85 931<br>5 806 / 1 841<br>108 |
| Bonds (rmsd)<br>Length (Å)<br>Angles (°) | <br>0.004<br>0.638 | <br>0.003<br>0.548 | <br>0.002<br>0.546 |
| *MolProbity* score | 2.57 | 2.01 | 1.98 |
| Clash score | 13.0 | 12.4 | 13.5 |
| Ramachandran plot (%)<br>Favoured<br>Allowed<br>Outliers | <br>92.35<br>7.36<br>0.29 | <br>94.84<br>5.01<br>0.16 | <br>95.16<br>4.68<br>0.16 |
| Rotamer outliers (%) | 3.92 | 2.72 | 3.40 |
| Cβ outliers (%) | 0.0 | NA | NA |
| Peptide plane (%)<br>Cis proline / general<br>Twisted proline / general | <br>1.8 / 0.0<br>0.0 / 0.0 | <br>2.1 / 0.0<br>0.0 / 0.0 | <br>2.2 / 0.0<br>0.0 / 0.0 |
| ADP B-factors (mean)<br>Protein<br>Nucleotides<br>Ligand | <br>89.19<br>73.00<br>54.85 | <br>82.07<br>64.48<br>62.43 | <br>54.32<br>64.93<br>44.42 |
| Resolution (Å)<br>used during refinement<br>$d_{99}$ (full/half1/half2)<br>$d_{model}$<br>$d_{FSC\text{-}model\ (0\ /\ 0.143\ /\ 0.5)}$ | masked / unmasked<br>2.8<br>3.1 / 3.2<br>3.0 / 3.0<br>2.6 / 2.8 / 2.9 //<br>2.7 / 2.8 / 3.1 | masked / unmasked<br>3.3<br>3.3 / 3.2<br>3.1 / 3.1<br>2.7 / 2.8 / 3.0 //<br>2.8 / 2.9 / 3.2 | masked / unmasked<br>3.4<br>3.5 / 3.5<br>3.4 / 3.4<br>2.9 / 3.0 / 3.3 //<br>3.0 / 3.2 / 3.7 |
| Model versus data<br>CC (mask)<br>CC (box)<br>CC (peaks)<br>CC (volume)<br>Mean CC for ligands | <br>0.81<br>0.72<br>0.66<br>0.80<br>0.76 | <br>0.82<br>0.72<br>0.66<br>0.81<br>0.72 | <br>0.78<br>0.72<br>0.66<br>0.76<br>0.77 |

**Table 2  Atomic model refinement and validation statistics**
